# Development of an orally bioavailable mSWI/SNF ATPase degrader and acquired mechanisms of resistance in prostate cancer

**DOI:** 10.1101/2024.02.29.582768

**Authors:** Tongchen He, Caleb Cheng, Yuanyuan Qiao, Hanbyul Cho, Eleanor Young, Rahul Mannan, Somnath Mahapatra, Stephanie J. Miner, Yang Zheng, NamHoon Kim, Victoria Z. Zeng, Jasmine P. Wisniewski, Siyu Hou, Bailey Jackson, Xuhong Cao, Fengyun Su, Rui Wang, Yu Chang, Bilash Kuila, Subhendu Mukherjee, Sandeep Dukare, Kiran B. Aithal, Samiulla D.S., Chandrasekhar Abbineni, Costas A. Lyssiotis, Abhijit Parolia, Lanbo Xiao, Arul M. Chinnaiyan

## Abstract

Mammalian switch/sucrose non-fermentable (mSWI/SNF) ATPase degraders have been shown to be effective in enhancer-driven cancers by functioning to impede oncogenic transcription factor chromatin accessibility. Here, we developed AU-24118, a first-in-class, orally bioavailable proteolysis targeting chimera (PROTAC) degrader of mSWI/SNF ATPases (SMARCA2 and SMARCA4) and PBRM1. AU-24118 demonstrated tumor regression in a model of castration-resistant prostate cancer (CRPC) which was further enhanced with combination enzalutamide treatment, a standard of care androgen receptor (AR) antagonist used in CRPC patients. Importantly, AU-24118 exhibited favorable pharmacokinetic profiles in preclinical analyses in mice and rats, and further toxicity testing in mice showed a favorable safety profile. As acquired resistance is common with targeted cancer therapeutics, experiments were designed to explore potential mechanisms of resistance that may arise with long-term mSWI/SNF ATPase PROTAC treatment. Prostate cancer cell lines exposed to long-term treatment with high doses of a mSWI/SNF ATPase degrader developed SMARCA4 bromodomain mutations and ABCB1 overexpression as acquired mechanisms of resistance. Intriguingly, while SMARCA4 mutations provided specific resistance to mSWI/SNF degraders, ABCB1 overexpression provided broader resistance to other potent PROTAC degraders targeting bromodomain-containing protein 4 (BRD4) and AR. The ABCB1 inhibitor, zosuquidar, reversed resistance to all three PROTAC degraders tested. Combined, these findings position mSWI/SNF degraders for clinical translation for patients with enhancer-driven cancers and define strategies to overcome resistance mechanisms that may arise.

**Significance Statement:** The mSWI/SNF complex is a promising therapeutic target for enhancer-driven cancers. PROTACs, which enable the targeting of “undruggable” proteins, often face the challenge of achieving oral bioavailability. Here, we present AU-24118, a first-in-class, orally bioavailable mSWI/SNF ATPase dual degrader with remarkable efficacy in *in vitro* and *in vivo* models. Additionally, our study describes two distinct mechanisms of resistance to PROTAC degraders, providing crucial insights into potential challenges facing their clinical application. These findings are critical for advancing PROTAC-based therapies to clinical settings as targeted therapies for cancers.

## Introduction

Epigenetic dysregulation has recently emerged as a promising avenue for innovative therapeutic approaches in the treatment of human cancer (1, 2). The mammalian switch/sucrose non-fermentable (mSWI/SNF) complex, a key player in this realm, exerts its influence on chromatin topology and gene expression by orchestrating the movement of nucleosome substrates (3). Comprising 11-15 subunits organized into canonical BAF (cBAF), polybromo-associated BAF (pBAF), and non-canonical BAF (ncBAF) forms, the complex is fueled by two ATPases, SMARCA2 and SMARCA4, which hydrolyze ATP to power the mSWI/SNF complexes (4–8). Recent research has underscored the potential of targeting these ATPase components, as perturbing them has been shown to impede tumor maintenance and oncogenicity (9–12). Notably, multiple small-molecule compounds specifically targeting ATPase mSWI/SNF subunits have been developed, with ongoing clinical trials (NCT04879017, NCT04891757) evaluating SMARCA2/4-targeting small molecules (13–16).

Concurrently, the field of cancer treatment research has witnessed a surge in interest in proteolysis targeting chimera (PROTAC) technology (17, 18). Several drugs based on PROTAC degraders are undergoing clinical trials, showcasing their unique ability to target previously deemed “undruggable” proteins, including transcription factors (19–21). We previously reported a highly selective PROTAC degrader targeting both mSWI/SNF ATPase subunits, SMARCA2 and SMARCA4, demonstrating remarkable therapeutic efficacy in various preclinical models of advanced prostate cancer through intravenous administration (16). Despite the success, the transition to oral dosing regimens for clinical applications becomes imperative for enhanced practicality and reduced invasiveness.

However, along with the rise in utilization of PROTAC degraders, a new challenge emerges: cancer cells develop resistance to PROTAC degraders over time (22, 23). Given the mechanism of action of PROTAC degraders, bringing into proximity the target of interest and an E3 ligase, mutations in any of these components can compromise the degradation system (24–26). Notably, long-term treatment may lead to mutations in the protein of interest, while the persistent expression of ATP binding cassette subfamily B member 1 (ABCB1), also known as multidrug resistance 1 (MDR1), poses an additional resistance mechanism observed in various therapies, including PROTAC treatment (22, 25, 27, 28).

Addressing these challenges, we developed a second-generation dual ATPase PROTAC degrader, AU-24118, which serves as a first-in-class, orally bioavailable degrader of the mSWI/SNF ATPase subunits. AU-24118 retains exquisite specificity for its targets while exhibiting increased potency in *in vitro* and *in vivo* models. Additionally, we characterized two independent potential resistance mechanisms to mSWI/SNF PROTAC degraders in prostate cancer models: one driven by mutations in the bromodomain of SMARCA4 and the other driven by ABCB1 elevation. Overall, this study leads the field in developing a novel, orally bioavailable mSWI/SNF ATPase PROTAC degrader and elucidates possible resistance mechanisms to these near-term clinical compounds in cancer, providing crucial technology and knowledge for future therapeutic development.

## Results

### Second-generation mSWI/SNF ATPase degrader AU-24118 exhibits potent and specific degradation of SMARCA2/4 and PBRM1

Previous studies by our group established that the degradation of mSWI/SNF ATPases disrupts enhancer-promoter looping interactions, resulting in a pronounced inhibition of tumor growth in xenograft models of prostate cancer (16). However, our first-generation compound, AU-15330, utilized the von Hippel-Lindau (VHL) E3 ligase, which likely contributed to its lack of oral bioavailability (**Fig. 1A**). Given the importance of developing an orally available drug for its practicality and cost-effectiveness, we generated AU-24118 as a first-in-class, second-generation mSWI/SNF ATPase degrader amenable to oral administration (**Fig. 1A**).

**Figure 1:**
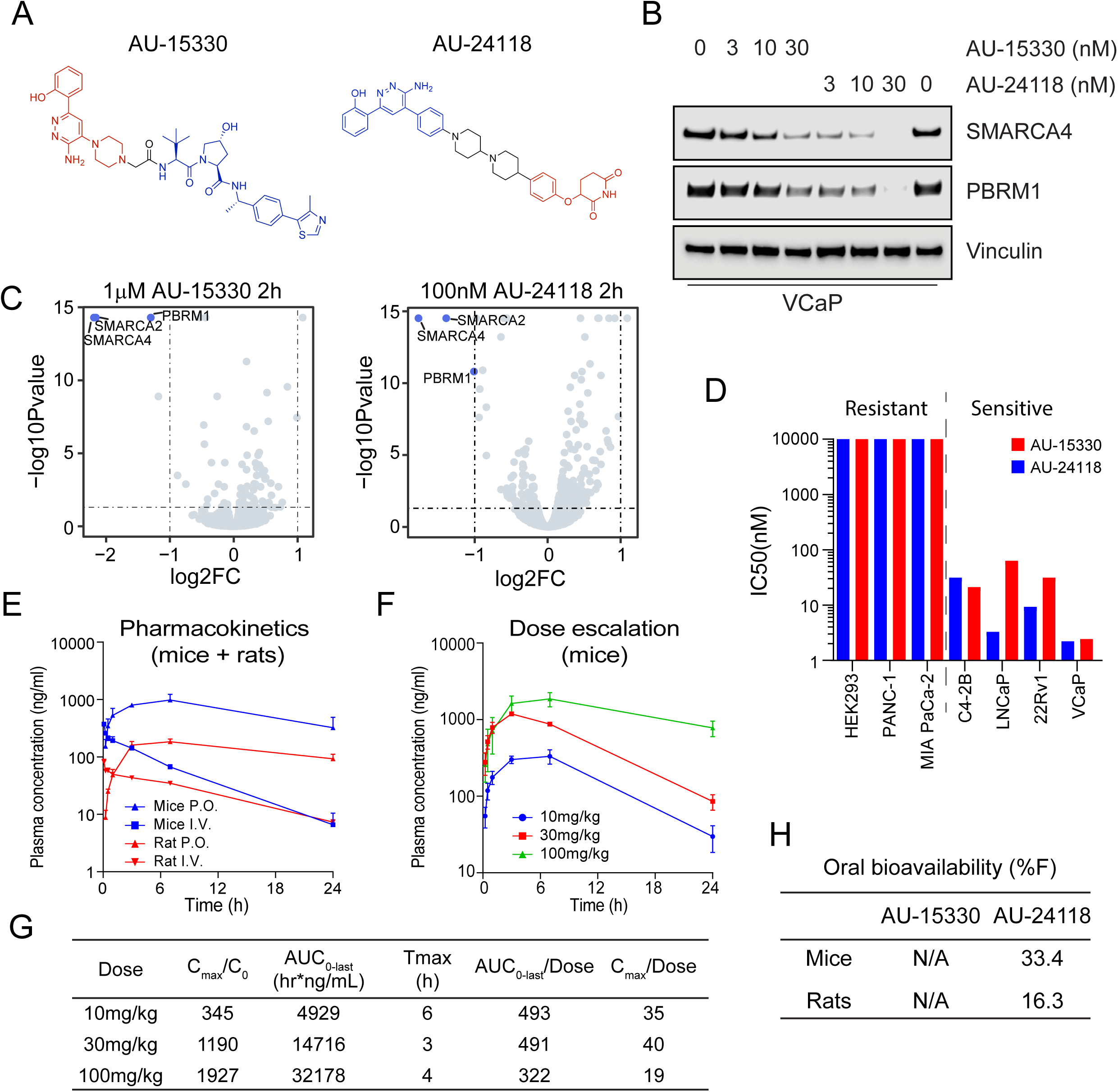
AU-24118 is a first-in-class, orally bioavailable degrader of the mSWI/SNF ATPases SMARCA2/4 and PBRM1. A. Chemical structures of first (AU-15330) and second (AU-24118) generation mSWI/SNF ATPase degraders. B. Immunoblot analysis of VCaP cells assessing protein levels of SMARCA4 (BRG1) and PBRM1 upon treatment with indicated doses of AU-15330 or AU-24118 for 4 hours. Vinculin was used as a loading control for all immunoblots unless otherwise indicated. C. Effects of AU-15330 and AU-24118 (2-hour treatment) on the proteome of VCaP cells. Data plotted are Log2 of the fold change (FC) versus DMSO control against -Log10 of the p-value from 3 biological replicates. T-tests were performed as two-tailed t-tests assuming equal variances. D. IC_50_ values of AU-15330 and AU-24118 in a panel of human cancer or normal cell lines after 5 days of treatment determined by Cell-TiterGlo viability assays. Cell lines are color-coded based on the treatment. E. Plasma concentrations over time of AU-24118 in mice or rats upon one-time oral (p.o.) or intravenous (i.v.) dose of AU-24118. F. Plasma concentrations over time of AU-24118 in mice upon oral dosing of indicated concentrations of AU-24118. G. Pharmacokinetics metrics for AU-24118 in mice as determined by dose-escalation analyses conducted according to panel F. Cmax = Peak Concentration, C0 = Initial Concentration, AUC_0-last_ = Area Under the Curve from Time 0 to Last Measurable Concentration, Tmax = Time to Peak Concentration. H. Oral bioavailability (%F) of AU-15330 and AU-24118 in mice and rats.

In accordance with the notion that cereblon (CRBN)-based PROTAC degraders tend to have higher oral bioavailability (29, 30), we chose to utilize CRBN as the E3 ligase for next-generation degraders. Several degraders were designed by conjugating ligands of SMARCA2/4 bromodomain and CRBN E3 ligase binders by employing knowledge-based iterative medicinal chemistry approaches. Further prioritization of designs for synthesis was made by utilizing an in-house proprietary ternary complex modeling algorithm, leading to the discovery of AU-24118 as a very potent, selective, and orally bioavailable degrader of SMARCA2, SMARCA4, and PBRM1 (**Fig. 1A**). Both compounds, AU-15330 and AU-24118, integrate a bait moiety binding to the bromodomains of SMARCA2 and SMARCA4, along with a ligand moiety for CRBN ligase (AU-24118) or VHL ligase (AU-15330) (see **SI Appendix, Fig. S1A**). In addition to targeting ATPases SMARCA2 and SMARCA4, both PROTAC degraders also exhibit binding to the secondary mSWI/SNF module component, PBRM1.

Under identical treatment conditions, AU-24118 demonstrated comparable degradation efficacy of SMARCA4 and PBRM1 to AU-15330 in VCaP prostate cancer cells (**Fig. 1B**). Further, to evaluate the specificity of AU-24118 to its target proteins, we employed mass spectrometry-based proteomics analysis, validating that AU-24118 selectively degraded SMARCA2, SMARCA4, and PBRM1, resulting in their significant downregulation following short-term treatment in a similar fashion to AU-15330 (**Fig. 1C**). Notably, this selectivity was maintained across all bromodomain-containing proteins and non-targeted mSWI/SNF subunits for AU-24118, similar to AU-15330, irrespective of the treatment duration (short-term (two hours) or long-term (eight hours)) (see **SI Appendix, Fig. S1B-E**).

Having determined that AU-24118 specifically degraded SMARCA2, SMARCA4, and PBRM1, we assessed the impact of AU-24118 and AU-15330 on cell viability across diverse cell lines. AU-24118 exhibited a notable preference for inhibiting transcription factor-addicted cancer cells, similar to AU-15330, as evidenced by half-maximal inhibitory concentrations (sensitive: IC_50_ < 100nM, resistant: IC_50_ > 100nM; **Fig. 1D**). To confirm the mechanism of action of AU-24118, specifically its utilization of the CRBN E3 ligase, the proteasome machinery, and the ubiquitination cascade, we co-treated VCaP cells with lenalidomide (CRBN binding ligand) or carfilzomib (proteasome inhibitor), which impeded AU-24118-induced degradation (see **SI Appendix, Fig. S1F**). In contrast, co-treatment with VL-285 (free VHL ligand) did not block target protein degradation. Taken together, these data demonstrate that AU-24118 potently and specifically degrades SMARCA2, SMARCA4, and PBRM1 through the CRBN-based proteasomal degradation machinery.

### AU-24118 is a CRBN-based, orally bioavailable PROTAC degrader of the ATPase subunits of the mSWI/SNF complex

To ascertain the oral bioavailability of AU-24118, we conducted pharmacokinetic analyses in mice and rats. Following oral gavage, plasma concentrations rose within 30 minutes and were maintained at steady states for over 24 hours (**Fig. 1E**). Subsequent dose escalation profiling at 10 mg/kg, 30 mg/kg, and 100 mg/kg revealed a concentration-dependent increase in plasma levels, demonstrating a dose-proportional elevation in area under the curve (AUC) (**Fig. 1F-G**). Notably, at 100 mg/kg, Cmax saturation, denoted by Cmax/dose, suggested potential absorption pathway saturation (**Fig. 1G**). Additionally, the detected free drug remained above the half-maximal effective concentration (EC_50_) for approximately 18 hours at the 30 mg/kg oral gavage dose (see **SI Appendix, Fig. S2A**). AU-24118 exhibited favorable pharmacokinetic properties with an overall oral bioavailability of 33.4% in mice (**Fig. 1H**, see **SI Appendix, Fig. S2B**).

To evaluate the preclinical safety of AU-24118, we conducted extensive toxicity analyses with AU-24118 in mice. First, to establish the maximum tolerated dose, we implemented a dosage escalation experiment in CB17 SCID tumor-free mice. Randomized into different arms and dosed three times weekly, mice receiving 20 mg/kg exhibited noticeable weight loss after the first week, culminating in morbidity or fatality by the second week. In contrast, all other dosage groups, notably including those receiving up to 15 mg/kg, demonstrated well-tolerated responses to AU-24118 (see **SI Appendix, Fig. S3A**).

We further characterized the host organ (spleen, kidney, and liver) morphology and function upon AU-24118 treatment at the maximum tolerated dose (15 mg/kg) in CB17 SCID mice. Hematoxylin-eosin (H&E) staining revealed no discernible toxicity in normal tissues (see **SI Appendix, Fig. S3B**) even upon successful target degradation as confirmed by immunohistochemistry (IHC) (see **SI Appendix, Fig. S3B**). Additionally, we evaluated liver and kidney function using a blood panel on a separate cohort of CB17 SCID mice that underwent 25-day treatment with AU-24118 at the maximum tolerated dose. Aside from a slight elevation in alanine transaminases (ALT), all other values were comparable to vehicle-treated mice and within normal ranges (see **SI Appendix, Fig. S3C**). This suggests that the loss of mSWI/SNF complex function had minimal impact on host organs compared to tumor tissue and that AU-24118 may have a favorable safety profile if translated to the clinic.

### AU-24118 treatment induces tumor regression in the castration-resistant prostate cancer (CRPC) VCaP xenograft model and is enhanced with concurrent enzalutamide treatment

Following dosage optimization and toxicity evaluations, we employed a CRPC xenograft model (CRPC-VCaP) to assess the therapeutic efficacy of AU-24118 (**Fig. 2A**). Post-castration and tumor regrowth, mice were randomly treated with vehicle, enzalutamide (10 mg/kg, oral gavage, five times weekly), AU-24118 (15 mg/kg, oral gavage, three times weekly), or both drugs simultaneously (**Fig. 2A**). Enzalutamide, an AR antagonist approved for treatment of CRPC, induced moderate tumor growth inhibition, as previously reported in this model (16). Intriguingly, AU-24118 treatment led to regression in every tumor, with the combination therapy demonstrating superior tumor regression, rendering some tumors unpalpable (**Fig. 2B-D**, see **SI Appendix, Fig. S3D**). Western blot analysis confirmed robust downregulation of direct targets (SMARCA2, SMARCA4, and PBRM1) and specific prostate cancer lineage proteins (AR, ERG, C-Myc, and PSA) after five days of AU-24118 treatment, whether administered alone or in combination with enzalutamide (**Fig. 2E**). Notably, AU-24118 treatment induced apoptosis as evidenced by increased cleaved PARP levels (**Fig. 2E**) as well as TUNEL staining (**Fig. 2F**). IHC confirmed SMARCA4 degradation and downregulation of AR, ERG, C-Myc, and the proliferation marker Ki-67 (**Fig. 2G**). Importantly, no significant body weight changes were observed throughout the treatments (see **SI Appendix, Fig. S3E**), indicating the mice tolerated treatment with AU-24118 with or without the addition of enzalutamide.

**Figure 2:**
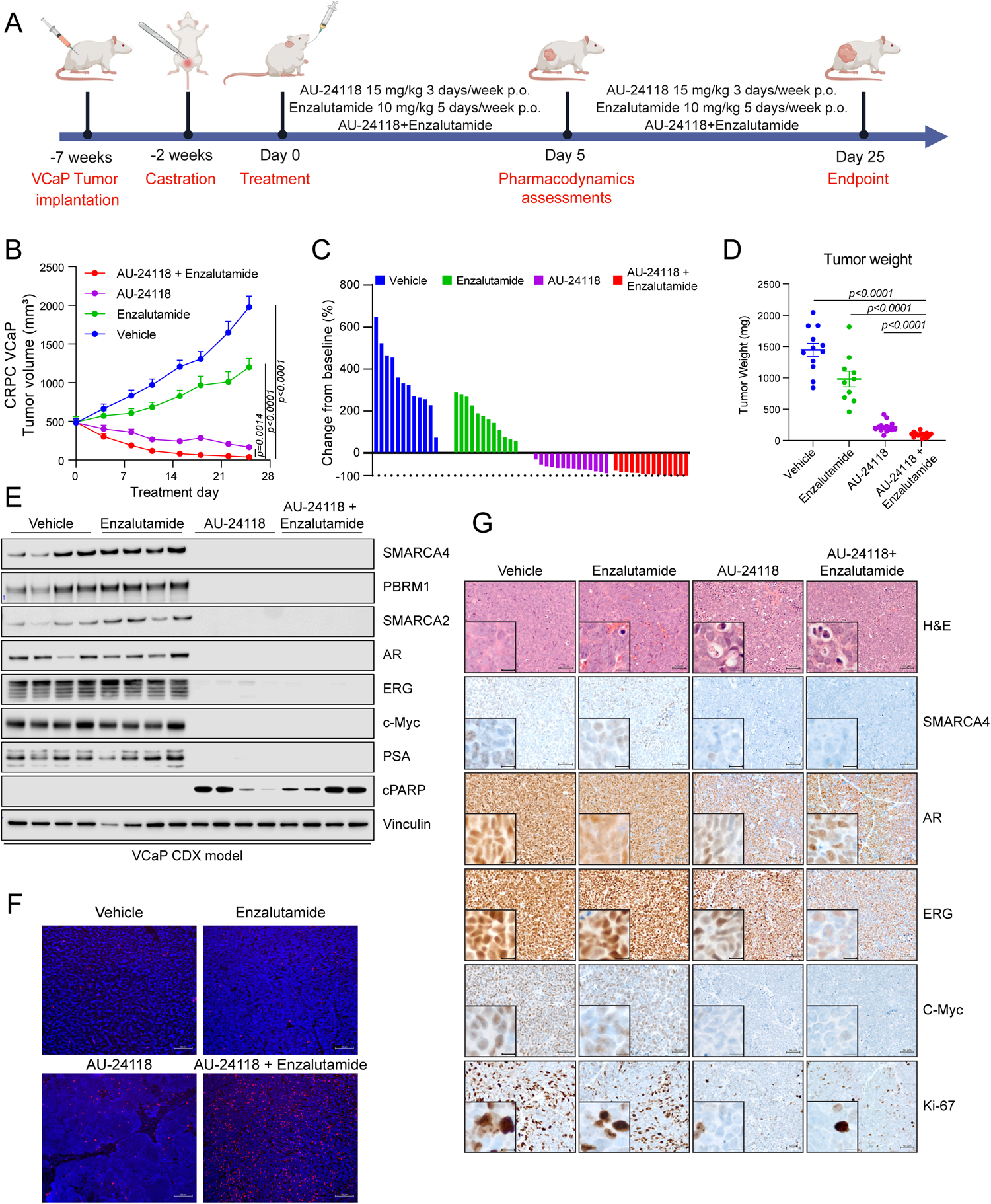
**AU-24118 induces tumor regression as a single agent, as well as in combination with enzalutamide, in a VCaP CRPC model.** A. Schematic of *in vivo* efficacy study of AU-24118 with or without enzalutamide in a VCaP castration-resistant prostate cancer (CRPC) model. VCaP-bearing mice were castrated and, upon tumor regrowth, randomized into treatment arms of vehicle, AU-24118, enzalutamide, or both AU-24118 and enzalutamide at indicated doses. B. Tumor measurements showing efficacy of AU-24118, enzalutamide, and AU-24118 + enzalutamide measured bi-weekly (analyzed with a two-way ANOVA with Dunnett’s multiple comparisons tests). Data shown are average tumor volumes +SEM. C. Waterfall plots showing change in tumor volume compared to baseline (day 0) at endpoint (day 25) of individual tumors of each treatment arm. D. Weights of individual tumors in each treatment arm at endpoint (day 25). Statistics were performed with a two-tailed unpaired t-test. E. Immunoblot of direct AU-24118 targets (SMARCA4, PBRM1, SMARCA2), validated downstream targets (AR, ERG, c-MYC, PSA/KLK3), and cleaved PARP (cPARP) in 4 tumors taken down at day 5 of treatment for pharmacodynamics assessment in the VCaP cell line-derived xenograft (CDX) model. F. Representative DAPI and TUNEL staining from VCaP xenograft of each treatment arm after 5 days treatment (scale=100 μm). G. Representative H&E staining with corresponding IHC analysis of direct AU-24118 target SMARCA4 and downstream targets (AR, ERG, c-MYC), and proliferation marker Ki-67 after 5 days of treatment (scale=50 μm). The inset scale=20 μm.

### Mutations in the PROTAC binding site of SMARCA4 confer resistance to mSWI/SNF ATPase degraders

The rising importance of the mSWI/SNF complex as a therapeutic target and the success (31, 32) of developing next-generation PROTAC degraders targeting the mSWI/SNF complex prompted us to proactively study potential resistance mechanisms that cancer cells may develop in response to this class of compounds. To achieve this, we cultured 22Rv1 cells in either DMSO or a lethal dose of AU-15330 for one month, refreshing the drug twice a week, until resistant cells emerged. Upon obtaining a population of resistant cells, we unbiasedly characterized the genomic and transcriptomic alterations using whole-exome sequencing and bulk RNA sequencing, respectively (**Fig. 3A**). Among the four independent resistant cell lines generated (AU-resistant, AUR), two were found to have point mutations in *SMARCA4* (lines AUR-1 and AUR-2), while the other two had wild-type *SMARCA4* (lines AUR-3 and AUR-4) (**Fig. 3B**). In contrast, the two lines (AUR-3/4) that did not harbor *SMARCA4* mutations were found to have high expression of ATP binding cassette subfamily B member 1 (*ABCB1*) while the other two (AUR-1/2) had low or no expression of *ABCB1* (**Fig. 3B**, see **SI Appendix, Fig. S4A**).

**Figure 3:**
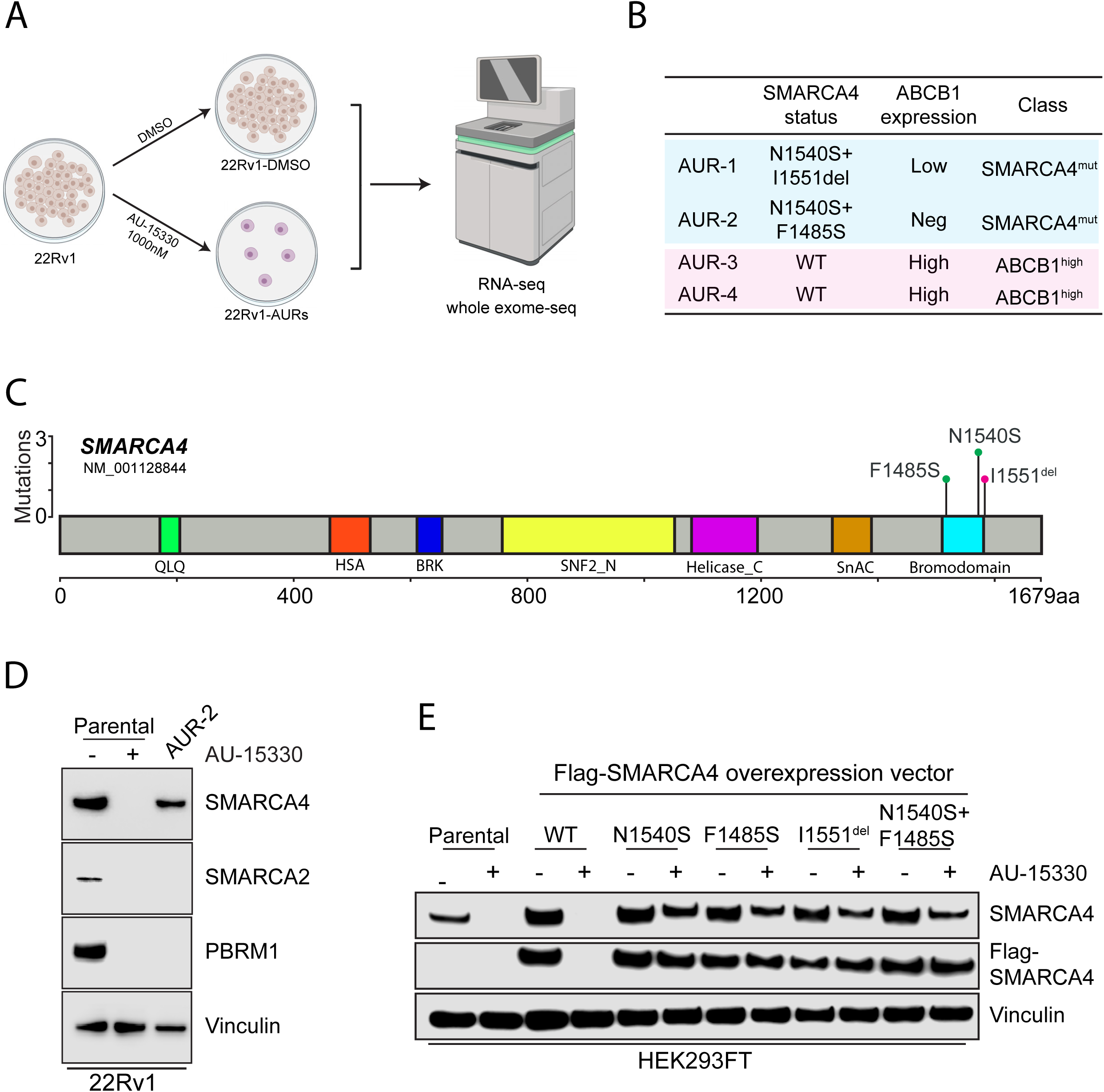
**Mutations in the SMARCA4 bromodomain as a mechanism of resistance to mSWI/SNF ATPase degraders.** A. Schematic of process used to generate and characterize four independent AU-15330-resistant (AUR) 22Rv1 cell lines. B. Table of 22Rv1 cell lines, highlighting SMARCA4 mutation profile, ABCB1 expression status, and classification of resistance mechanism. C. Lollipop plot of SMARCA4 mutations detected through whole-exome sequencing of AUR-1 and AUR-2 cell lines. Amino acid residues are plotted on the X axis with protein domains highlighted and labeled. Mutation frequencies are plotted on the Y axis. D. Immunoblot analysis of the indicated target proteins in 22Rv1 wild-type (WT) or 22Rv1 AUR-2 cells cultured with 1 μM AU-15330. E. Immunoblot of HEK293 cells transfected with labeled overexpression vectors or untransfected (parental) for 48 hours prior to treatment with AU-15330 (1μM, +) or DMSO (-) for 24 hours showing changes in SMARCA4 or FLAG (FLAG-SMARCA4).

To characterize the *SMARCA4* mutations in AUR-1/2 cells, we first mapped the point mutations of the two lines and determined that they all occurred in or near the bromodomain of SMARCA4, the binding site of both AU-15330 and AU-24118 (**Fig. 3C**). To determine the functional relevance of the SMARCA4 mutations, we compared SMARCA4, SMARCA2, and PBRM1 levels of AUR-2 cells grown in AU-15330 (1 μM) to wild-type 22Rv1 cells acutely treated with AU-15330 (1 μM) or DMSO. As expected, SMARCA4, SMARCA2, and PBRM1 levels were eliminated by AU-15330 in wild-type cells; however, only SMARCA2 and PBRM1 levels were eliminated in AUR-2 cells by AU-15330, while SMARCA4 was maintained at a detectable level (**Fig. 3D**). Similarly, even though AU-15330 and AU-24118 were able to degrade SMARCA2 and PBRM1, they were unable to degrade SMARCA4 in AUR-2 cells (see **SI Appendix, Fig. S4B**) or cause a decrease in cell viability (see **SI Appendix, Fig. S4C**). Finally, to further determine the functional role of the SMARCA4 mutations found in AUR-1/2 cells, we cloned overexpression vectors of flag-tagged wild-type or mutant SMARCA4. In accordance with our hypothesis, overexpressed mutant forms of SMARCA4 in HEK293 were resistant to degradation by AU-15330 (**Fig. 3E**). Taken together, mutations in or near the bromodomain of SMARCA4 that prevent degradation of SMARCA4 are sufficient to allow for prostate cancer cell survival in the presence of mSWI/SNF ATPase degraders.

### Sustained degradation of mSWI/SNF ATPases may induce overexpression of ABCB1 as a resistance mechanism

To characterize the potential resistance mechanisms of AUR-3/4 cell lines which did not carry the SMARCA4 mutations, we conducted transcriptomics analyses on AUR-3/4 cells. Through RNA sequencing, we identified that *ABCB1* expression, which is basally undetectable in wild-type 22Rv1 cells, was highly upregulated (**Fig. 4A**). We further confirmed this expression at the protein level and determined that ABCB1 expression was not modulated with acute AU-15330 treatment in AUR-3 cells (**Fig. 4B**). To ascertain whether ABCB1 overexpression could be observed in *in vivo* settings, we analyzed tumor samples from the endpoints of multiple efficacy studies featuring AU-15330 detailed in our previous study (16). In a C4-2B prostate cancer cell line-derived xenograft model described in our previous study (16), we detected *ABCB1* overexpression at the transcript (**Fig. 4C**) and protein (**Fig. 4D**, see **SI Appendix, Fig. S5A**) levels in the tumors at endpoint (day 24), but not at day five (see **SI Appendix, Fig. S5B-C**). Additionally, in the MDA-146-12 prostate cancer patient-derived xenograft model, we found an upregulation of *ABCB1* after 43 days of treatment with AU-15330 (see **SI Appendix, Fig. S5D**). Taken together, these data suggest that *ABCB1* can be upregulated upon long-term treatment with AU-15330.

**Figure 4:**
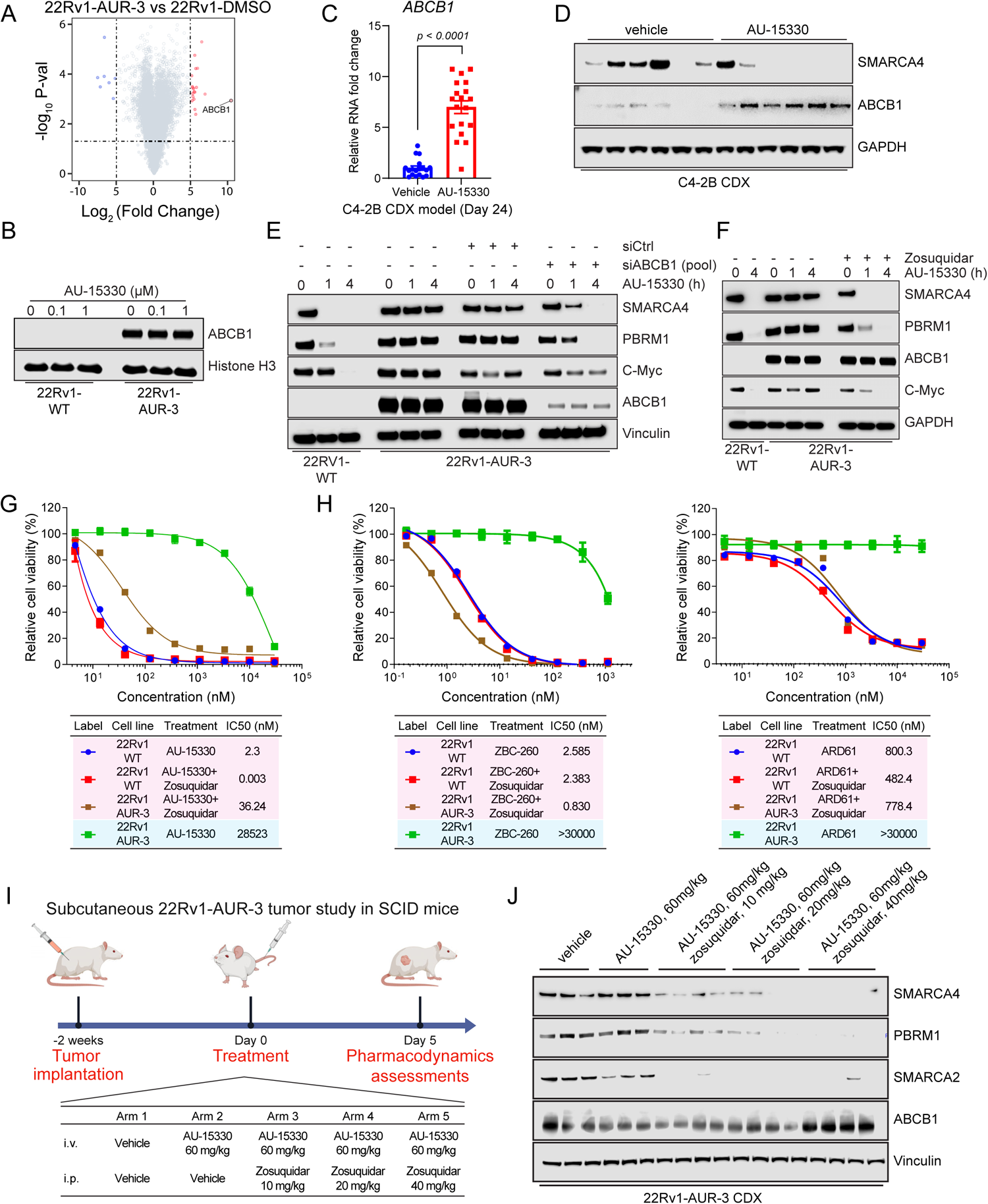
**ABCB1/MDR1 overexpression as a mechanism of resistance to PROTAC degraders which can be overcome with ABCB1 inhibition.** A. Volcano plot visualizing the overall transcriptomic alterations as assessed by RNA-seq in 22Rv1-AUR-3 versus 22Rv1-WT cells. B. Immunoblot analysis of 22Rv1-wild type (WT) or 22Rv1 AUR-3 cells treated with DMSO, 0.1 μM AU-15330, or 1 μM AU-15330 for 4 hours. Histone H3 was used as a loading control. C. qPCR analysis of C4-2B cell line-derived xenograft (CDX) tumors upon 24 days of dosing with either vehicle or AU-15330 showing changes in mRNA level of *ABCB1*. T-tests were performed as two-tailed t-tests assuming equal variances. D. Immunoblot illustrating changes in protein level of ABCB1 and SMARCA4 in C4-2B CDX tumors after long-term AU-15330 treatment. Vinculin is utilized as the loading control across immunoblots. E. Immunoblot analysis of 22Rv1-wild type (WT) or 22Rv1 AUR-3 cells treated with DMSO, si-ABCB1 (pooled), or si-control (siCtrl, pooled) and 1μM AU-15330 for the indicated durations. Vinculin is utilized as the loading control across immunoblots. F. Immunoblot analysis of 22Rv1-wild type (WT) or 22Rv1 AUR-3 cells treated with DMSO, 1 μM zosuquidar, and 1 μM AU-15330 for the indicated timepoints. Vinculin is utilized as the loading control across immunoblots. G. Dose-response curves of 22Rv1-wild type (WT) or 22Rv1 AUR-3 cells treated with AU-15330 with or without zosuquidar (1 μM). Data are presented as mean +/− SD (n = 6) from one-of-three independent experiments. IC_50_ values calculated from dose-response curve experiment. H. (Left) Dose-response curves of 22Rv1-wild type (WT) or 22Rv1 AUR-3 cells treated with ZBC-260 with or without zosuquidar (1 μM). Data are presented as mean +/− SD (n = 6) from one-of-three independent experiments. IC_50_ values calculated from dose-response curve experiment. (Right) Dose-response curves of 22Rv1-wild type (WT) or 22Rv1 AUR-3 cells treated with ARD-61 with or without zosuquidar (1 μM). Data are presented as mean +/− SD (n = 6) from one-of-three independent experiments. IC_50_ values calculated from dose-response curve experiment. I. Schematic of *in vivo* pharmacodynamics study of AU-15330 with indicated doses of zosuquidar and AU-15330 in a 22Rv1-AUR-3 xenograft model. J. Immunoblot illustrating levels of the indicated proteins in 22Rv1-AUR-3 xenografts after 5-day treatment with the indicated dosing regimens. Vinculin is utilized as the loading control across immunoblots.

As ABCB1 confers resistance through transport of small molecules out of the cell, ABCB1 as a resistance mechanism should theoretically be compound-agnostic. Thus, we tested whether AUR-3 cells would also be resistant to other PROTAC degraders. Indeed, AUR-3 cells were dramatically less sensitive to bromodomain-containing protein 4 (BRD4) degrader ZBC-260 and androgen receptor (AR) degrader ARD-61 (see **SI Appendix, Fig. S5E**). Knowing that ABCB1 confers broad resistance to these other PROTAC degraders, we sought to determine whether chronic treatment with other PROTAC degraders could also induce ABCB1 upregulation. Thus, following the previously described procedure for AU-15330 (**Fig. 3A**), we chronically treated 22Rv1 cells with ZBC-260 at a lethal dose. After four weeks of treatment, we found that resistant cells indeed induced *ABCB1* overexpression (see **SI Appendix, Fig. S5F**). Given this, we concluded that ABCB1 could be induced upon long-term treatment with PROTAC degraders and that it provides broad resistance to many PROTAC degraders.

To evaluate the functional relevance of ABCB1, we knocked down ABCB1 in AUR-3 cells and showed that ABCB1 knockdown partially restored the efficacy of AU-15330 in degrading its target proteins (**Fig. 4E**) and decreasing viability in AUR-3 cells (see **SI Appendix, Fig. S5G**). Likewise, we determined that zosuquidar, an ABCB1 inhibitor (33), restored sensitivity of AUR-3 cells to AU-15330 (**Fig. 4F-G**). Accordingly, co-treatment with zosuquidar also restored sensitivity of AUR-3 cells to the other PROTAC degraders (ZBC-260, ARD-61) (**Fig. 4H**, see **SI Appendix, Fig. S5H**), directly implicating ABCB1 in providing resistance to various types of PROTAC degraders.

Given the translational relevance of mSWI/SNF ATPase degraders for potential cancer therapeutics, we sought to determine whether ABCB1-dependent resistance mechanisms to PROTAC degraders could be modulated *in vivo*. To evaluate this, we established a 22Rv1-AUR-3 cell line-derived xenograft model and assessed the ability of zosuquidar to sensitize 22Rv1-AUR-3 tumors to AU-15330. Indeed, upon 5 days of treatment, AU-15330 was unable to degrade any of its target proteins alone; however, simultaneous dosing of zosuquidar restored the degradation efficacy of AU-15330 in a dose-dependent manner (**Fig. 4I-J**). This suggests that ABCB1-dependent resistance mechanisms to PROTAC degraders can be attenuated by concurrent pharmacologic inhibition of ABCB1.

## Discussion

PROTAC technology has expanded the possibilities of targeted cancer therapies by overcoming a multitude of challenges that traditional small-molecules have faced (34). For instance, PROTAC degraders allow for targeting of traditionally difficult to inhibit proteins. This is due to their mechanism of bringing the target and an E3 ubiquitin ligase in proximity, which allows for target perturbation without necessitating binding to a specific active site pocket (35). Furthermore, PROTAC degraders can be administered at reduced concentrations, extended dosing intervals, and with lower toxicity in comparison to inhibitors of the same targets. This is because their effective protein degradation occurs at low concentrations, and they are not limited by equilibrium occupancy. Administering lower doses of PROTACs may decrease the likelihood of off-target effects occurring (36, 37). Encouragingly, more than 20 PROTAC degraders have already undergone various stages of clinical testing. For example, ARV-110 (AR degrader) and ARV-471 (estrogen receptor degrader) are currently in phase II/III clinical trials. This indicates that PROTACs have the potential to be well-tolerated therapeutics with potent capabilities to modulate cancer-specific targets.

Our laboratory previously reported on AU-15330, a first-in-class PROTAC degrader of the mSWI/SNF ATPases SMARCA2 and SMARCA4. We further showed that degradation of these subunits, along with PBRM1, led to rapid and widespread chromatin compaction, perturbing essential oncogenic programs driven by transcription factors such as AR, FOXA1, and ERG in prostate cancer models. Phenotypically, we showed that transcription factor-driven cancer models were preferentially sensitive to mSWI/SNF ATPase degradation while non-transcription factor-driven cancer or normal models were relatively insensitive. This work laid the foundation for further clinical development of therapeutics targeted toward the mSWI/SNF ATPases to strategically treat transcription factor-driven cancers. Specifically, development of an orally bioavailable mSWI/SNF ATPase degrader to improve practicality of drug administration and cost was pursued, leading to AU-24118.

In addition to the numerous benefits of being orally bioavailable, AU-24118 also has increased efficacy compared to AU-15330 in terms of both target degradation efficiency as well as potency *in vivo*. For example, in many of our *in vitro* assays, we utilized roughly a 10-fold lower concentration of AU-24118 than AU-15330 to achieve comparable effects. Additionally, in the VCaP CRPC model described in **Fig. 2**, we found that AU-24118 (15 mg/kg) as a single agent regressed every tumor in the cohort whereas AU-15330 (60 mg/kg) had more modest effects as a single agent.

Paralleling the future of PROTAC degraders as precision therapies for cancers will inevitably be a rise in resistance mechanisms to such compounds. In our study, we created numerous independent AU-15330-resistant prostate cancer lines and identified two classes of resistance mechanisms: one by which mutations in the SMARCA4 binding site of the drug is mutated, and the other by which ABCB1 is overexpressed. Resistant lines harboring mutations in SMARCA4 were completely insensitive to the mSWI/SNF ATPase degraders both in terms of SMARCA4 degradation as well as any cellular phenotypic effects. However, given that both AU-15330 and AU-24118 have a similar binding site, cells harboring this mutation may still be sensitive to independent SMARCA4 inhibitors/degraders with unique binding sites. This finding highlights the importance of mutational identification through sequencing to identify potentially resistant cancers prior to enrolling patients in a treatment regimen featuring PROTAC degraders.

An additional implication of the SMARCA4 mutations being a mechanism of acquired resistance to mSWI/SNF ATPase degraders is that SMARCA4 may be critical while SMARCA2 and PBRM1 are more dispensable in the presence of functional SMARCA4 (38, 39). In two independently generated AU-15330-resistant cell lines, three unique mutations in SMARCA4 were found, but none were found in SMARCA2 or PBRM1. Indeed, AU-15330-resistant cells cultured in the presence of AU-15330 had sustained loss of SMARCA2 and PBRM1 while retaining wild-type levels of SMARCA4. This implies that SMARCA4, rather than SMARCA2 or PBRM1, may be the critical target to perturb in the context of prostate cancer therapy.

ATP binding cassette subfamily B member 1 (ABCB1/P-gp/MDR1) has been identified as a resistance mechanism to various therapeutics, including chemotherapies, kinase inhibitors, and PROTAC degraders (40, 41). Here, we find that ABCB1 is also a mechanism of acquired resistance to mSWI/SNF ATPase degraders that confer broad resistance to multiple PROTAC degraders. Genetic or pharmacological inhibition of ABCB1 (using zosuquidar) reversed the resistance and restored PROTAC efficacy. We specifically utilized zosuquidar in this study as it is a known ABCB1 inhibitor that has been tested in a phase III clinical trial and, thus, has a favorable safety profile (42). However, many ABCB1 inhibitors, including compounds directed toward ABCB1 as well as those that can be repurposed to inhibit ABCB1, have been described (43, 44). For example, important recent work has demonstrated additional promising avenues to attenuate ABCB1-driven resistance mechanisms to PROTAC degraders such as using the EGFR/ErbB inhibitor, lapatinib (22). The development of safer and more potent compounds to ABCB1 will be critical for successful clinical application of PROTAC degraders for cancer therapy.

Our present study describes a major step toward translating mSWI/SNF ATPase PROTAC degraders to clinical use through developing a first-in-class, orally bioavailable compound and demonstrating its preclinical safety and efficacy. Additionally, we explore possible resistance mechanisms of prostate cancer models to long-term treatment with mSWI/SNF ATPase degraders and nominate potential strategies to mitigate them. Taken together, our data emphasizes the promise of PROTAC degraders targeting epigenetic regulators while highlighting potential future challenges of implementing such compounds as cancer therapies.

## Methods

### Cell lines, antibodies, and compounds

Every cell line utilized in this study was originally sourced from the American Type Culture Collection (ATCC). Cells were cultured at 37 °C in an atmosphere of 5% CO_2_. Cell lines were profiled using short tandem repeats (STR) to confirm their identity at the University of Michigan (U-M) Sequencing Core. They were also tested every two weeks for the presence of mycoplasma. LNCaP and 22Rv1 cells were grown in Gibco RPMI-1640 + 10% FBS (Thermo Fisher). VCaP cells were grown in Gibco DMEM + 10% FBS (Thermo Fisher). The sources of all antibodies are described in the **SI Appendix, Table S1**. AU-15330 and AU-24118 were synthesized by Aurigene (see **SI Appendix, Supplementary Methods**). Zosuquidar was purchased from ApexBio. ARD-61 and ZBC-260 were generously provided by Dr. Shaomeng Wang (U-M).

### Cell viability assays

Cells were seeded onto 96-well plates in appropriate culture media and allowed to adhere overnight prior to treatment. The next day, a 9-point serial dilution of the compounds was prepared and added to the plates. After treatment, the cells were then incubated for an additional five days. On the last day of treatment, the CellTiter-Glo cell viability assay (Promega) was then executed in accordance with the manufacturer’s guidelines. The signal (luminescence) from each well was determined using an Infinite M1000 Pro plate reader (Tecan). Relative viability was determined by dividing the signal from each well by the average of the control condition in each experiment. Finally, the data were analyzed and plotted using GraphPad Prism.

### Immunoblots

Lysates were prepared using RIPA buffer (Thermo Fisher) with cOmpleteTM protease inhibitor cocktail tablets (Sigma-Aldrich) added. Total protein for each sample was determined using the DC Protein Assay (Bio-Rad) according to the manufacturer’s instructions. A standardized quantity of protein was resolved using NuPAGE Tris-Acetate Protein Gel (3-8%)(Thermo Fisher) or NuPAGE Bis-Tris Protein Gel (4-12%) (Thermo Fisher) and probed with primary antibodies as indicated. Following incubation with secondary antibodies conjugated with HRP, membranes were imaged on an Odyssey Fc Imager (LiCOR Biosciences).

### RNA isolation and quantitative real-time PCR

Total RNA was extracted from cell samples using the miRNeasy mini kit (Qiagen). Complementary DNA (cDNA) was synthesized from 1000 ng of RNA using the Maxima First Strand cDNA Synthesis Kit for PCR with reverse transcription (RT–PCR) (Thermo Fisher). Quantitative PCR (qPCR) was performed using standard SYBR green reagents and protocols on a QuantStudio 7 Real-Time PCR system (Applied Biosystems). All reactions were performed in technical triplicates. The expression of the target mRNA was determined using the ΔΔCt method using beta-Actin expression as a reference for each sample. All the primers utilized were designed using Primer 3 (http://frodo.wi.mit.edu/primer3/) and synthesized by Integrated DNA Technologies. Primer sequences are listed in the **SI Appendix, Table S2**.

### RNA-seq and analysis

RNA-seq libraries were prepared using 800 ng of total RNA by KAPA RNA HyperPrep RiboErase and mRNA kits. Ribosomal RNA was depleted, poly(A) mRNA was selected by Sera-Mag™ Magnetic Oligo(dt) particles (GE38152103011150) respectively, and then fragmented to around 200-300bp with heat in fragmentation buffer. cDNA synthesis, end-repair, A-tailing, and ligation of the NEB adapters were all performed following the KAPA RNA Hyper protocols (KAPA RNA HyperPrep Kit with RiboErase (HMR) and KAPA mRNA HyperPrep kit, Roche). Libraries were size-selected for 300– 500 bp fragments by double AMPure beads- and PCR-amplified using 2x KAPA HiFi HotStart mix and NEB dual indexes. Library quality (concentration and product size) was measured on an Agilent 2100 Bioanalyzer (DNA 1000 chip). Paired-end libraries were sequenced with the Illumina NovaSeq 6000, (paired-end 2 × 151 nucleotide read length) with sequence coverage of 30-40 million paired reads. and data analysis was performed as previously described (16).

### Genomic DNA extraction and whole-exome sequencing

Upon isolating total DNA from cells using the DNeasy Blood & Tissue Kit (Qiagen), 1–3 μg of genomic DNA was sheared using a Covaris ME220 to a peak target size of 300 base pairs (bp). Fragmented DNA subsequently underwent end-repair, followed by A-tailing, ligated with NEB adapters-, and size-selected with double AMPure beads. Fragments with sizes in the range of 350 to 550 bp were recovered, amplified with 2x KAPA HiFi HotStart mix and NEB dual indexes, and purified using AMPure beads. 1 μg of the library was hybridized to the Agilent SureSelect XT Human All Exon V4 probes (Agilent Technologies 5190-4634). The targeted exon fragments were captured by Streptavidin beads (Fisher 65602), washed, and enriched following Agilent’s protocol. The whole-exome library quality was measured on an Agilent 2100 Bioanalyzer (DNA 1000 chip) for concentration and product size. Paired-end libraries were sequenced with the Illumina NovaSeq 6000, (paired-end 2 × 151 nucleotide read length) with sequence coverage of 100-150 million paired reads.

Novoalign was used to align the FASTQ data from whole-exome libraries to the GRCh38 reference genome. Data were then transformed into BAM files using SAMtools. The primary somatic call-set was generated using Strelka based on the following criteria: allele frequency exceeding 0.05 in the 22Rv1-AURs, allele frequency less than 0.01 in the normalized data, a minimum of five variant reads, normal depth surpassing 50, and Somatic Evidence Score (EVS) over the 90th percentile of the overall EVS distribution. These detected signals were further enhanced by mutations identified by VarDict in genes that are repeatedly mutated in prostate cell lines 22RV1. Visual representation of mutation calls in *SMARCA4* (NM_00128844) was facilitated using cBioPortal.

### SMARCA4mut plasmid construction and transfection

GFP-SMARCA4 (Addgene, 65391) was used for the SMARCA4 mutation clone. The pLVX-TetOne-Puro-hAXL (Addgene, 124797) backbone was used as a starting point prior to undergoing a series of modifications. The endogenous target gene was removed, and a multiple cloning site (MCS) was inserted downstream of the endogenous promoter. Custom gblocks containing Flag-tagged wild-type SMARCA4 were synthesized (IDT) and inserted into the MCS. Subsequently, custom gblocks containing SMARCA4-N1540S, SMARCA4-I1551del, or SMARCA4-F1485S were inserted to replace the respective portion of wild-type SMARCA4. Finally, the TRE3G promoter was replaced with hPGK. HEK293 cells were seeded overnight and transfected with Lipofectamine 3000 (Thermo Fisher) according to the manufacturer’s protocol. Media was changed eight hours post transfection and incubated for 48 hours prior to conducting further analyses.

### siRNA

Cells were seeded in a 6-well plate. After overnight incubation, cells were then transfected with 25 nM of either ON-TARGETplus Human ABCB1 (5243) SMARTpool or non-targeting pool siRNAs (negative control, Dharmacon) using the Lipofectamine RNAiMAX reagents (Thermo Fisher). Following 24 hours of incubation, cells were trypsinized and re-seeded into 6 well plates. After another overnight incubation, cells underwent an additional round of transfection following the same protocol. Following 48 hours of incubation, cells were used for further experimentation.

### TMT mass spectrometry

Cells were plated at 5 × 10^6^ cells per well in a 6-well plate overnight prior to treatment with DMSO, 100 nM AU-24118, or 1 μM AU-15330 for 2 hours. Whole cell lysates were prepared in RIPA buffer (Thermo Fisher) supplemented with tablets of cOmpleteTM protease inhibitor cocktail (Sigma-Aldrich). For each condition, 75 μg of total protein was labelled with TMT isobaric Label Reagent (Thermo Fisher) according to the manufacturer’s protocol and subjected to 12 fractions of liquid chromatography–mass spectrometry (LC–MS)/MS analysis.

### AU-24118, enzalutamide, AU-15330, and zosuquidar formulation for *in vivo* studies

AU-15330 was mixed into 40% 2-hydroxypropyl-β-cyclodextrin (HPβCD) and sonicated until the powder was completely dissolved. This solution was then mixed with 5% dextrose in water (D5W) to achieve a final concentration of 10% HPβCD. AU-15330 was delivered to mice by intravenous injection through retro-orbital injection. AU-24118 was mixed into PEG200 and then sonicated and vortexed until completely dissolved. Five volumes of 10% D-α-Tocopherol polyethylene glycol 1000 succinate were then added and vigorously mixed. Four volumes of 1% Tween-80 were then added followed by another round of vigorous shaking. AU-15330 and AU-24118 were freshly prepared immediately prior to administration to mice. Enzalutamide was added in 1% carboxymethyl cellulose (CMC) with 0.25% Tween-80, followed by sonication until homogenized. AU-24118 and enzalutamide were delivered to mice by oral gavage. Zosuquidar was dissolved in DMSO, and then the solution was further mixed with nine volumes of 10% HPβCD. Zosuquidar was delivered to mice by intraperitoneal injection.

### Human tumor xenograft models

Six-week-old CB17 (male) severe combined immunodeficiency (SCID) mice were obtained from the breeding colony at the University of Michigan. For all studies, tumors were measured two or three times per week using digital calipers, and tumor volumes were calculated using the formula (π/6) (L × W2), where L is length and W is width of the tumor. At study endpoint, mice were sacrificed, and tumors were extracted and weighed. For the VCaP castration-resistant tumor model, 3 × 10^6^ VCaP cells were subcutaneously injected into the dorsal flank on both sides of each mouse in a 1:1 mixture of serum free medium and Matrigel (BD Biosciences). Once tumors grew to a palpable size (∼400 mm^3^), the mice were physically castrated. Once those tumors regained their pre-castration size, mice were then randomized and treated with either 15 mg/kg AU-24118 or vehicle by oral gavage 3 days per week with or without enzalutamide (10 mg/kg) by oral gavage 5 days per week for 4 weeks.

For the 22Rv1-AUR-3 model, bilateral subcutaneous dorsal flank tumors were established by injecting 1 × 10^6^ cells suspended in a 1:1 mixture of serum-free medium and Matrigel (BD Biosciences). Once tumors reached an average volume of 100 mm^3^, mice were randomized and treated with either 60 mg/kg AU-15330 by retro-orbital vein injection 3 times every other day and with or without 10 mg/kg, 20 mg/kg, or 40 mg/kg of zosuquidar by intraperitoneal injection daily for five days per week. In accordance with the Institutional Animal Care and Use Committee (IACUC) guidelines, the maximal tumor size did not exceed the 2.0 cm limit in any dimension for any treatment condition. Animals harboring tumors approaching that size were immediately euthanized. All *in vivo* studies were approved by the U-M IACUC.

### Drug toxicity assessments using histopathological analysis of organs

Organs (liver, spleen, and kidney) were harvested from mice and fixed in neutral buffered formalin (10%) and subsequently embedded in paraffin to generate tissue blocks. These blocks were then sectioned at a thickness of 4 µm and stained using Harris haematoxylin (Leica Surgipath) and alcoholic eosin-Y stain (Leica Surgipath). Staining was executed using the automated Leica autostainer-XL platform. Upon staining, the sections were subsequently analyzed by two pathologists on a brightfield microscope to determine morphology and architectural coherence. These pathological assessments were performed in a blinded manner. Detailed and comprehensive analysis of cellular and -subcellular level changes of organs were performed as previously described (16).

### Immunohistochemistry

Immunohistochemistry (IHC) was performed on 4 µm formalin-fixed, paraffin-embedded tissue sections using BRG1 (SMARCA4), AR, FOXA1, ERG and C-Myc. The IHC process was performed on the Ventana ULTRA automated slide staining system using the Omnimap and Ultraview Universal DAB detection kit. The antibody details are provided in the **SI Appendix, Table S1**. The following commercial kits from Roche-Ventana Medical System were used: Discovery CC1 (Cat No. 950-500), Discovery CC2 (Cat No. 950-123), OptiView Universal DAB Detection Kit (Cat No. 760-700), and OmniMap Univesral DAB Detection Kit (Cat No. 760-149).

### *In vivo* apoptosis evaluation

The Terminal dUTP Nick End Labeling (TUNEL) and In Situ Cell Death Detection Kits (TMR Red #12156792910; Roche Applied Science) were used to examine apoptosis, following the manufacturer’s instructions. Briefly, fixed sections were permeabilized with Triton X-100 and then washed in PBS. Labelling was carried out at 37°C for 60 minutes by addition of a reaction buffer that contained enzymes. A Zeiss Axiolmager M1 microscope was used to obtain images.

### *In vivo* pharmacokinetics studies

All experimental procedures relating to animal care, handling, and treatment were approved by the Institutional Animal Ethics Committee (IAEC) of Aurigene Oncology Ltd. based on the Committee for Control and Supervision of Experiments on Animals (CCSEA) guidelines. Animals were included for experimentation after one week of acclimatization to standard laboratory conditions and were provided with RO-treated water and rodent pellet diet ad libitum.

The pharmacokinetic profile of AU-24118 was studied in both mice and rats. For assessment of oral bioavailability, AU-24118 was administered by intravenous (i.v.) and oral (p.o.) routes at 1 mg/kg and 30 mg/kg doses, respectively (n = 3). For the oral arm, AU-24118 was administered as a solution formulation in 10% PEG200, 50% of 20% D-α-Tocopherol polyethylene glycol 1000 succinate and 40% of 1% Tween-80 in water while AU-24118 was formulated in 2% DMA and 10% HPβCD in saline (qs) for the intravenous administration.

### Determination of plasma drug concentrations

Concentrations of AU-24118 were determined in plasma samples using a non-validated (fit for purpose) LC-MS/MS method in positive Multiple Reaction Monitoring (MRM) mode with transition pair of m/z: 633.11→344.00. Either Carbamazepine (with a transition pair of m/z: 237.07→194.10) or Telmisartan (with a transition pair of m/z: 515.16→276.00) were used as internal standards. Sample processing was carried out by protein precipitation followed by evaporation method. Compound concentrations were calculated using Analyst® software 1.7.2 with linear/quadratic regression algorithm (with 1/x2 weighting factor) over a quantification range of 1 ng/mL to 2500 ng/mL with the low and high ends of this range defining the lower limit of quantitation (LLOQ) and upper limit of quantitation (ULOQ), respectively.

### Statistical analysis

Independent samples were used for the acquisition of all data points. The analysis and visualization of data were performed using Prism version 9 (GraphPad Software; San Diego, CA). Results were expressed as means ± standard deviation (SD) or ± standard error of the mean (SEM), as indicated in the figure legends. Statistical significance was established with a p-value below 0.05, unless otherwise indicated.

## Data availability

RNA-sequencing and whole exome-sequencing data have been deposited in the NCBI Gene Expression Omnibus (GEO) with the accession number GSE250328. All other data are available in the main text or the supplementary information.

## Acknowledgements

We express our gratitude to Alex Hopkins, Lisa McMurry, Amanda Miller, Christine Caldwell-Smith, Xia Jiang, Heng Zheng, Sarah Yee, Yunhui Cheng, Rupam Bhattacharyya, Jianzhou Xu, Chia-Mei Huang, Melissa Dunn, Jillian Schell, and Shuqin Li from the Michigan Center for Translational Pathology at the University of Michigan, as well as Venkatesha Basrur and the Rogel Cancer Center Proteomics Shared Resource. We also thank Susanta Samajdar and Murali Ramachandra from Aurigene Oncology Limited for their guidance. This research received primary funding from the Trailsend Foundation, with additional support from the Prostate Cancer Foundation (PCF), National Cancer Institute (NCI) Prostate Specialized Programs of Research Excellence (SPORE) Grant P50-CA186786, and an NCI Outstanding Investigator Award R35-CA231996 (A.M.C.). C.C. is supported by the National Institutes of Health Cellular and Molecular Biology Training Grant (5T32-GM145470). L.X. is a recipient of the 2022 PCF Young Investigator Award, and he also benefits from support provided by the Department of Defense Prostate Cancer Research Program Idea Development Award (W81XWH-21-1-0500) and the Michigan SPORE Career Enhancement Program. A.M.C. is a Howard Hughes Medical Institute Investigator, A. Alfred Taubman Scholar, and American Cancer Society Professor.

## Author Contributions

T.H., C.C., L.X., and A.M.C. conceived the study, designed the experiments, and composed the manuscript. A.M.C. and L.X. supervised the project. Y.Q. assisted with study conception and design. T.H., C.C., and L.X. performed all *in vitro* experiments with assistance from N.H.K., V.Z., Y.C., and J.P.W. H.C., E.Y., and A.P. carried out all bioinformatic analyses with assistance from S.H. R.M. and S.Mahapatra carried out all histopathological evaluations and performed immunohistochemistry with assistance from Y.Z., N.H.K., and B.J. F.S., R.W., and X.C. generated next-generation sequencing libraries and performed the sequencing. S.J.M. and C.A.L. assisted with manuscript writing and organization. C.A., B.K., S.Mukherjee, S.D., K.B.A., and S.D.S. discovered AU-15330 and AU-24118 and carried out the PK/PD studies described in this study.

## Classification

Biological Sciences - Medical Sciences

