## Supplemental Information (Methods, Figures, and Tables) for "Development of an orally bioavailable mSWI/SNF ATPase degrader and acquired mechanisms of resistance in prostate cancer"

[1]Michigan Center for Translational Pathology, University of Michigan, Ann Arbor, MI, USA
[2]Department of Pathology, University of Michigan, Ann Arbor, MI, USA
[3]Department of Urology, Xiangya Hospital, Central South University, Changsha, Hunan, China
[4]Medical Scientist Training Program, University of Michigan, Ann Arbor, MI, USA
[5]Cellular and Molecular Biology Program, University of Michigan, Ann Arbor, MI, USA
[6] Rogel Cancer Center, University of Michigan, Ann Arbor, MI, USA
[7]Department of Biostatistics, School of Public Health, University of Michigan, Ann Arbor, MI, USA
[8]Howard Hughes Medical Institute, University of Michigan, Ann Arbor, MI, USA
[9]Aurigene Oncology Limited, Bangalore, India
[10]Department of Molecular and Integrative Physiology, University of Michigan, Ann Arbor, MI, USA
[11]Department of Internal Medicine, Division of Gastroenterology, University of Michigan, Ann Arbor, MI, USA
[12]Department of Urology, University of Michigan, Ann Arbor, MI, USA
[13]These authors contributed equally

**This PDF includes:**

Supplementary Methods
Figures S1 to S5
Tables S1 to S2

### Supplementary Methods

#### Synthesis of AU-24118

General Information: All chemicals and solvents were obtained from commercial suppliers and sed without further purification. Purification was performed using combi-flash Nextgen300. All reactions were monitored by TLC, using silica gel plates with fluorescence F254 and UV light visualization. $^1$H NMR spectra was recorded on a Varian Mercury Plus at 400 MHz, and $^{13}$C NMR spectra was recorded on a JEOL-ECZ-400S spectrometer at 100 MHz. Coupling constants (J) are expressed in hertz (Hz). Chemical shifts ( δ ) of NMR are reported in parts per million (ppm) units relative to internal control (TMS). Signal splitting patterns are described as singlet (s), doublet (d), triplet (t), quartet (q), multiplet (m), broad (br), or a combination thereof. The low resolution of ESI-MS was recorded on an Agilent-6120, and the high-resolution mass (resolution-70000) for the compound was generated using Q-Exactive Plus orbitrap system (Thermo Scientific) using electrospray ionization (ESI). Melting point was recorded in Stuart instrument, model smp30. HPLC was recorded in Agilent 1260 infinity II with PDA detector; the column used was a KINETEX EVO C-18(150mm x 4.6mm, 5 μ ) using A: 0.01%TFA IN WATER B: ACN 100%; Method -T/%B: 0/5,1/5,6/100,8/100,10/5,12/5 method and Flow rate: 1.0 ml/min. FTIR spectrum was recorded using Spectrum Two™ instrument (Scanning range 4000 cm-1 to 450 cm-1, Pallet making -For solid sample with KBr).

Abbreviations used: DMSO for dimethylsulfoxide, DIPEA for N, N-diisopropylethylamine, MeOH for methanol, DMF for N, N-dimethylformamide, THF for tetrahydrofuran, DCM for dichloromethane, NaH for Sodium hydride, KOAc for Potassium acetate, $K_2CO_3$ for Potassium carbonate, CuI for Copper(I) iodide, AcOH for acetic acid, HATU for 1- [bis(dimethylamino) methylene]-1H-1,2,3-triazolo[4,5-b]pyridinium 3-oxid hexafluorophosphate, Pd(dppf)$Cl_2$ for [1,1′ -bis(diphenylphosphino)ferrocene]dichloro-palladium(II).

**Chemistry Synthesis**

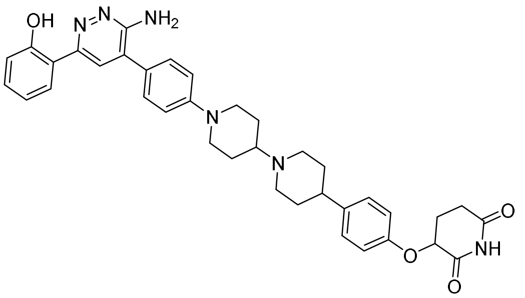

**Synthesis of 3-(4-(1'-(4-(3-amino-6-(2-hydroxyphenyl)pyridazin-4-yl)phenyl)-[1,4'-bipiperidin]-4-yl)phenoxy)piperidine-2,6-dione hydrochloride (AU-24118)**

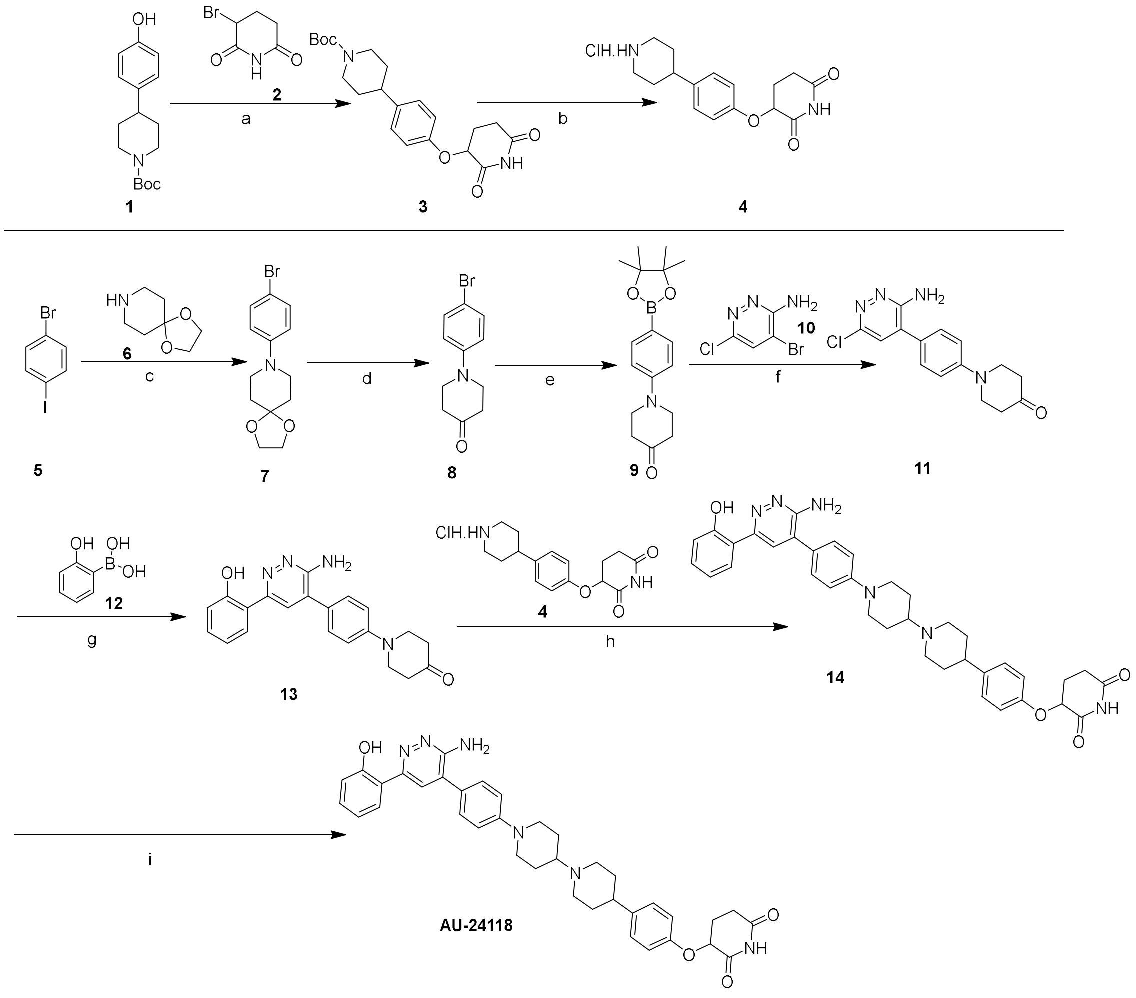


Reagents and conditions: a) NaH, DMF, 0 °C - RT, 2 h; b) 4M HCl in 1,4-dioxane, DCM, 0 °C - RT, 4h; c) $K_2CO_3$, CuI, L-proline DMF, 90 °C, 16h; d) 1M HCl (aq), THF, 80 °C, 6h; e) bis(pinacolato)diboron, Pd(dppf)$Cl_2$•DCM, KOAc, dioxane, sealed tube, 100 °C, 6 h; f) Pd(dppf)$Cl_2$•DCM, $K_2CO_3$, dioxane, water, sealed tube, 100 °C, 16 h; g) Pd(dppf)$Cl_2$•DCM,

$K_2CO_3$, dioxane, water, sealed tube, 120 °C, 4h; h) KOAc, AcOH, molecular sieves (4 A°), $NaBH(OAc)_3$, DMSO, THF, RT, 16 h; i) 4M HCl in 1,4-dioxane, DCM, 0 °C - RT, 1h.

Step-a: tert-butyl 4-(4-((2,6-dioxopiperidin-3-yl)oxy)phenyl)piperidine-1-carboxylate

To a stirred solution of tert-butyl 4-(4-hydroxyphenyl)piperidine-1-carboxylate (2.0g, 7.21 mmol) in DMF (20 mL) was added NaH (0.43g, 18.02 mmol) at 0°C temperature and stirred for 30 mins. Then 3-bromopiperidine-2,6-dione (1.66 g, 8.65 mmol) was added in the reaction mixture and stirred at RT for 2h. After completion of the reaction, the reaction mixture was poured in ice cold water and extracted with 2 X 100 mL of ethyl acetate, evaporated. The crude product was purified first by combi flash column chromatography using 20-30% Ethyl acetate in hexane to afford the title compound colourless solid (1.1g, 39 %). $^1$H NMR (400 MHz, DMSO-$d_6$): δ 10.91 (s, 1H), 7.14 (d, *J* = 8.8 Hz, 2H), 6.93 (d, *J* = 8.4 Hz, 2H), 5.15-5.13 (m, 1H), 4.10-3.90 (m, 2H), 2.84-2.72 (m, 2H), 2.68-2.55 (m, 2H), 2.20-2.11 (m, 3H), 1.73-1.69 (m, 2H), 1.51-1.42 (m, 2H), 1.41 (s, 9H). LCMS: m/z 387.1 (M-H).

Step-b: 3-(4-(piperidin-4-yl)phenoxy)piperidine-2,6-dione hydrochloride

To a stirred solution of tert-butyl 4-(4-((2,6-dioxopiperidin-3-yl)oxy)phenyl)piperidine-1-carboxylate (1.1g, 2.83 mmol) in DCM (10 mL) was added 4 N dioxane hydrochloride (10 mL) at 0 °C and then slowly brought to RT and stirred for 4h. The reaction mixture was evaporated under reduced pressure. The resultant residue was washed with diethyl ether and dried under vacuum to afford the title compound yellowish solid (0.81g, crude). $^1$H NMR (400 MHz, DMSO-$d_6$): δ 10.93 (s, 1H), 8.91 (bs, 1H), 8.71 (bs, 1H), 7.13 (d, *J* = 8.8 Hz, 2H), 6.97 (d, *J* = 8.4 Hz, 2H), 5.18-5.15 (m, 1H), 3.34-3.31 (m, 2H), 2.98-2.93 (m, 2H), 2.78-2.68 (m, 2H), 2.63-2.61 (m, 1H), 2.17-2.12 (m, 2H), 1.91-1.87 (m, 2H), 1.85-1.78 (m, 2H). LCMS: m/z 289.1 (M+H).

Step-c: 8-(4-bromophenyl)-1,4-dioxa-8-azaspiro[4.5]decane

To a stirred solution of 1-Bromo-4-iodobenzene (10g, 35.3 mmol), 1,4-dioxa-8-azaspiro[4.5]decane (5.06 g, 35.3 mmol) in DMSO (100 mL) were added $K_2CO_3$ (9.77g, 70.69 mmol) and L-proline (1.62g, 14.1 mmol), and the reaction mixture was degassed with nitrogen for 10 min. This was followed by CuI (1.34g, 7.06 mmol) added to the reaction mixture, and the reaction mixture was heated at 90 °C for 16 h in a sealed tube. Once the reaction was completed (monitored by TLC), the reaction mixture was diluted with EtOAc. The combined organic layer was washed with water, brine, dried over anhydrous sodium sulphate, and concentrated under vacuum to give the residue which was purified by combi flash column chromatography using 10% ethyl acetate in hexane as eluent to afford the title compound as off-white solid (7g, 66%). $^1$H NMR (400 MHz, DMSO-$d_6$): δ 7.31 (d, $J$ = 9.2 Hz, 2H), 6.90 (d, $J$ = 9.2 Hz, 2H), 3.90 (s, 4H), 3.27-3.24 (m, 4H), 1.68-1.65 (m, 4H); LC-MS: m/z 297.9 (M+H).

Step-d: 1-(4-bromophenyl)piperidin-4-one

To a stirred solution of 8-(4-bromophenyl)-1,4-dioxa-8-azaspiro[4.5]decane (7g, 13.41 mmol) in THF (50 mL) was added 1(N) aq. HCl (50 mL) at 0 ºC and then slowly brought to RT; the reaction mixture was heated for 6 h at 80 ºC. Once the reaction was completed (monitored by TLC), the reaction mixture was basified with $NaHCO_3$ and extracted with EtOAc. The combined organic layer was washed with water, brine, dried over anhydrous sodium sulphate, and concentrated under vacuum to give the residue which was purified by combi flash column chromatography using 10% ethyl acetate in hexane as eluent to afford the title compound as off-white solid (4g, 66%). $^1$H NMR (400 MHz, DMSO-$d_6$): δ 7.35 (d, $J$ = 8.8 Hz, 2H), 6.97 (d, $J$ = 8.8 Hz, 2H), 3.60-3.57 (m, 4H), 2.40-2.37 (m, 4H); LC-MS: m/z 254 (M+H).

Step-e: 1-(4-(4,4,5,5-tetramethyl-1,3,2-dioxaborolan-2-yl)phenyl)piperidin-4-one

To a stirred solution of 1-(4-bromophenyl)piperidin-4-one (4g, 15.74 mmol) in dioxane (80 mL) were added bis(pinacolato)diboron (5.99 g, 23.60 mmol) and KOAc (4.63g, 47.22 mmol) at RT and the reaction mixture was degassed with nitrogen for 10 min then $Pd(dppf)_2Cl_2$.DCM (1.28 g, 1.57 mmol) was added into the reaction mixture and the reaction mixture was heated at 100 ºC for 16 h. Once the reaction was completed (monitored by TLC), the reaction mixture was diluted with EtOAc. The combined organic layer was washed with water, brine, dried over anhydrous sodium sulphate and concentrated under vacuum to give the residue which was purified by combi flash column chromatography using 15% ethyl acetate in hexane as eluent to afford the title compound as off-white solid (4g, 84 %). $^1$H NMR (400 MHz, DMSO-$d_6$): δ 7.53 (d, $J$ = 8.4 Hz, 2H), 6.97 (d, $J$ = 8.4 Hz, 2H), 3.68-3.65 (m, 4H), 2.40-2.37 (m, 4H), 1.26 (s, 12 H); LC-MS: m/z 302.05 (M+H).

Step-f: 1-(4-(3-amino-6-chloropyridazin-4-yl)phenyl)piperidin-4-one

To a stirred solution of 4-bromo-6-chloropyridazin-3-amine (8.3g, 39.84 mmol), 1-(4-(4,4,5,5-tetramethyl-1,3,2-dioxaborolan-2-yl)phenyl)piperidin-4-one (10.0g, 33.20 mmol) in 1,4-dioxane (80 mL) and water (10 mL) was added $K_2CO_3$ (13.76g, 99.6 mmol) and degassed with nitrogen for 10 min. This was followed by adding Pd(dppf)$_2$Cl$_2$.DCM (2.71g, 3.32 mmol), and the reaction mixture was heated at 100 ºC in a sealed tube for 16 h. Once the reaction was completed (monitored by TLC), the reaction mixture was diluted with EtOAc. The combined organic layer was washed with water, brine, dried over anhydrous sodium sulphate, and concentrated under vacuum to get crude product which was purified by combi flash column chromatography using 70-90% EtOAc in hexane as eluent to afford the title compound white solid (5.0g, 50%). $^1$H NMR (400 MHz, DMSO-d$_6$): δ 7.47 (d, *J* = 8.8 Hz, 2H), 7.29 (s, 1H), 7.13 (d, *J* = 9.2 Hz, 2H), 6.28 (bs, 2H), 3.71 (t, *J* = 6.0 Hz, 4H), 2.43 (t, *J* = 6.4 Hz, 4H); LC-MS: m/z 302.95 (M+H).

Step-g: 1-(4-(3-amino-6-(2-hydroxyphenyl)pyridazin-4-yl)phenyl)piperidin-4-one

To a stirred solution of 1-(4-(3-amino-6-chloropyridazin-4-yl)phenyl)piperidin-4-one (11g, 36.33 mmol), (2-hydroxyphenyl)boronic acid (7.01g, 50.86 mmol) in 1,4-dioxane (50 mL) and water (13 mL) was added $K_2CO_3$ (15.0g, 138.2 mmol) and degassed with nitrogen for 10 min. This was followed by adding Pd(dppf)$_2$Cl$_2$.DCM (2.96 g, 3.63 mmol), and the reaction mixture was heated for 4 h at 120 ºC in a sealed tube. Once the reaction was completed (monitored by TLC), the reaction mixture was diluted with EtOAc. The combined organic layer was washed with water, brine, dried over anhydrous sodium sulphate, and concentrated under vacuum to give the residue which was purified by combi flash column chromatography using 90 - 100% EtOAc in hexane as eluent to afford the title compound off white solid (7.0g, 54%). $^1$H NMR (400 MHz, DMSO-d$_6$): δ 13.7 (s, 1H), 7.92 (m, 2H), 7.53 (d, *J* = 8.8 Hz, 2H), 7.21 (t, *J* = 7.6 Hz, 1H), 7.14 (d, *J* = 8.4 Hz, 2H), 6.89-6.83 (m, 2H), 6.36 (bs, 2H), 3.73 (t, *J* = 5.6 Hz, 4H), 2.45 (t, *J* = 5.6 Hz, 4H). LC-MS: m/z 361.0 (M+H).

Step-h: 3-(4-(1'-(4-(3-amino-6-(2-hydroxyphenyl)pyridazin-4-yl)phenyl)-[1,4'-bipiperidin]-4-yl)phenoxy)piperidine-2,6-dione

To a stirred solution of 3-(4-(piperidin-4-yl)phenoxy)piperidine-2,6-dione (0.500g, 1.73 mmol) and 1-(4-(3-amino-6-(2-hydroxyphenyl)pyridazin-4-yl)phenyl)piperidin-4-one hydrochloride (0.625g, 1.73 mmol) in 12 mL THF:DMSO (2:1) mixture was added KOAc (0.511g, 5.20 mmol) and acetic acid (0.2 mL) and molecular sieves (4Aº), and the reaction mixture was stirred at 70 ºC for 16h. Then the reaction mixture was cooled to 0ºC, and sodium triacetoxy borohydride (1.10 g, 5.20 mmol) was added; the reaction mixture was stirred at RT for 12h. The reaction was monitored by TLC. After completion of the reaction, the reaction mixture was extracted in DCM, and a saturated ammonium chloride wash was given to the organic layer followed by a saturated sodium chloride wash. The organic layer was dried over anhydrous sodium sulphate and then concentrated under reduced pressure, and the resultant residue was washed with diethyl ether and dried under vacuum to afford the title compound light yellow solid (0.310g, 28.25 %). $^{1}$H NMR (400 MHz, DMSO-$d_6$): δ 13.76 (s, 1H), 10.90 (s, 1H), 7.94 (d, *J* = 7.2 Hz, 1H), 7.93(s, 1H), 7.50 (d, *J* = 8.8 Hz, 2H), 7.23 (dt, *J* = 8.4, 1.6 Hz, 1H), 7.14 (d, *J* = 8.8 Hz, 2H), 7.07 (d, *J* = 8.8 Hz, 2H), 6.92-6.85 (m, 4H), 6.37 (bs, 2H), 5.15-5.11 (m, 1H), 3.89 (d, *J* = 12.4 Hz, 2H), 2.98 (d, *J* = 10 Hz, 2H), 2.82-2.55 (m, 4H), 2.48-2.35 (m, 2H), 2.30-2.05 (m, 4H), 1.87-1.82 (m, 2H), 1.76-1.67 (m, 2H), 1.61-1.48 (m, 4H). LCMS: m/z 633.05 (M+H).

<u>Step-i: 3-(4-(1'-(4-(3-amino-6-(2-hydroxyphenyl)pyridazin-4-yl)phenyl)-[1,4'-bipiperidin]-4-yl)phenoxy)piperidine-2,6-dione hydrochloride</u>

To a stirred solution of 3-(4-(1'-(4-(3-amino-6-(2-hydroxyphenyl)pyridazin-4-yl)phenyl)-[1,4'-bipiperidin]-4-yl)phenoxy)piperidine-2,6-dione (0.250g, 0.395 mmol) in DCM (2 mL) was added 4 N dioxane hydrochloride (2 mL) at 0 ºC and then slowly brought to RT and stirred for 1h. The reaction mixture was evaporated under reduced pressure. The resultant residue was washed with diethyl ether and dried under vacuum to afford the title compound yellowish solid (0.240 g, 94.6%). m.p.: 117-120ºC; HPLC purity 97.49% (RT- 4.75 min); $^{1}$H NMR (400 MHz, DMSO-$d_6$): δ 11.01 (bs, 1H), 10.89 (s, 1H), 8.10 (bs, 1H), 8.06 (s, 1H), 7.59 (dd, *J* = 1.2, 7.6 Hz, 1H), 7.50 (d, *J* = 8.8 Hz, 2H), 7.33 (t, *J* = 7.2 Hz, 1H), 7.17-7.11 (m, 4H), 7.05 (d, *J* = 7.6 Hz, 1H), 6.95 (d, *J* = 8.8Hz, 3H), 5.16-5.11 (m, 1H), 4.05 (d, *J* = 12.0 Hz, 2H), 3.51 (d, *J* = 11.2 Hz, 2H), 3.46-3.34 (m, 1H), 3.10-2.98 (m, 2H), 2.89-2.53 (m, 5H), 2.27-2.05 (m, 6H), 1.95-1.72 (m, 4H); 13C NMR (100 MHz, DMSO-d6) δ 174.2, 173.1, 171.9, 156.8, 156.1, 154.9 151.0, 150.4 137.6, 134.7, 133.4, 132.3, 130.1, 127.8, 122.2, 120.0, 117.2, 116.5, 116.0, 72.9, 62.6, 49.2, 47.4, 38.6, 30.3, 30.2, 25.5, 24.6 ; IR (KBr): 3405.82, 1705.00, 1654.94, 1605.89, 1510.97, 1459.52, 1245.90, 1196.03, 1069.34, 1004.54, 832.79, 765.40, 552.83 cm−1; HRMS (ESI) found 633.3176.

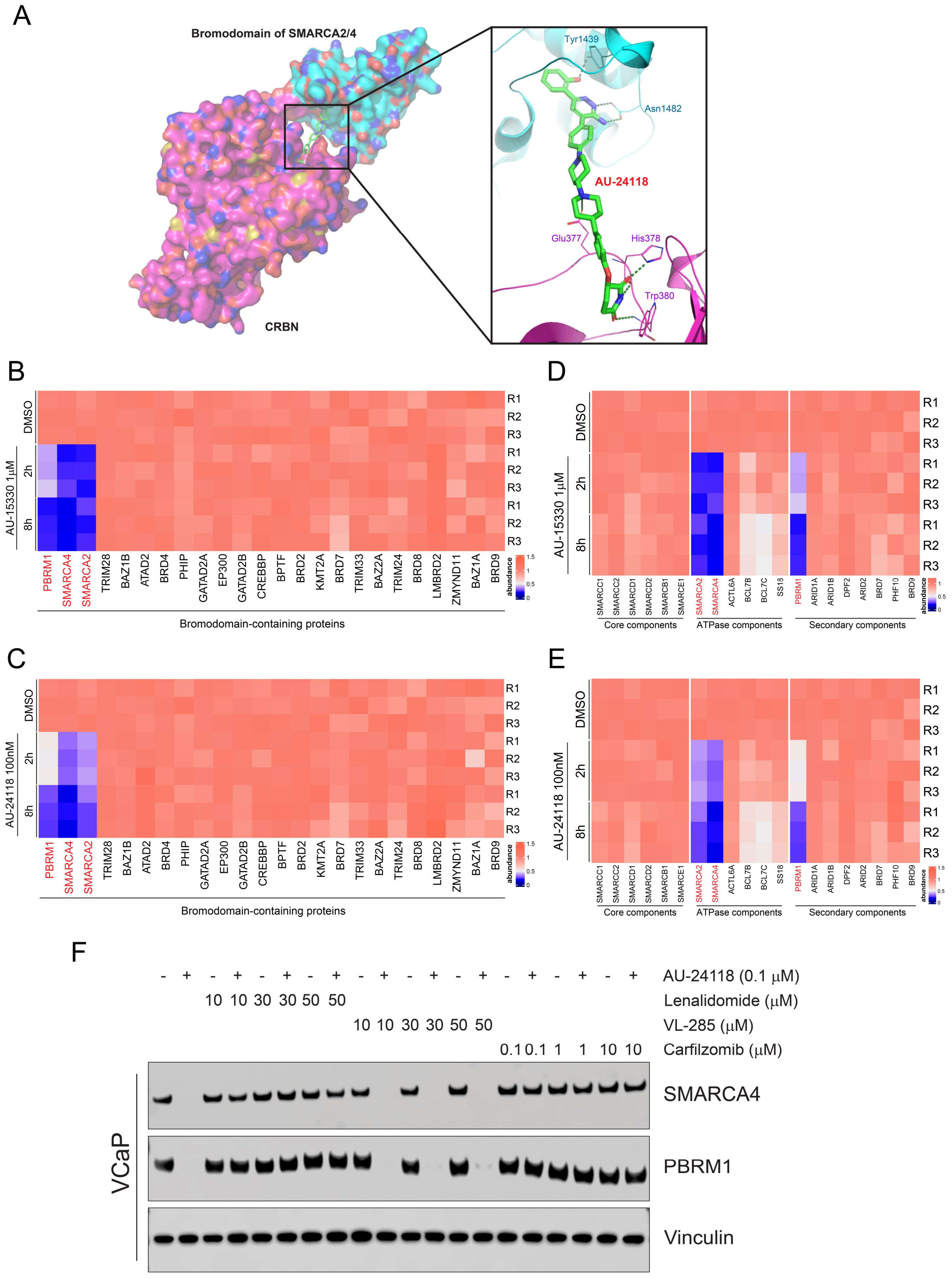


**Fig. S1. (A)** Docking model of AU-24118 (green sticks) with the SMARCA2/4 and CRBN complex. AU-24118 is suggested to fit into the pocket of SMARCA2/4 and CRBN and capture several key interactions. Key hydrogen bond interactions with protein residues (cyan sticks in SMARCA2/4, pink sticks in CRBN) are shown. **(B)** Heatmap of relative abundance of bromodomain-containing proteins detected via Tandem Mass Tag (TMT)-based quantitative MS upon 2-hour (2h) and 8-hour (8h) AU-15330 treatment (1μM). DMSO, dimethyl sulfoxide (vehicle). Data from three independent replicates are shown in panels B-E. **(C)** Heatmap of relative abundance of bromodomain-containing proteins detected via TMT-based quantitative MS upon 2-hour and 8 hour AU-24118 treatment (100 nM). **(D)** Heatmap showing TMT-based MS abundance of detectable mSWI/SNF components after 2 hours and 8 hours of treatment with AU-15330 (1μM). **(E)** Heatmap showing TMT-based MS abundance of detectable mSWI/SNF components after 2 hours and 8 hours of treatment with AU-24118 (100 nM). **(F)** Immunoblots of SMARCA4 and PBRM1 in VCaP cells pre-treated with VL285, lenalidomide, or carfilzomib for 1h and subsequently treated with AU-24118 at indicated concentrations for 4h. Vinculin was used as a loading control on all immunoblots. This experiment was repeated independently twice.

A

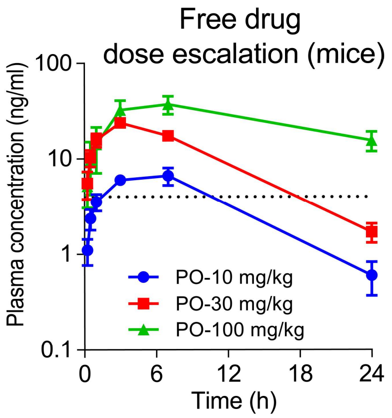


B

| PK parameters | Units | CD1 Mice | | Wistar Rat | |
|---|---|---|---|---|---|
| | | I.V. | P.O. | I.V. | P.O. |
| $AUC_{0\text{-}INF}$ | hr*ng/mL | 1675 | NC | 756 | NC |
| $AUC_{0\text{-}last}$ | hr*ng/mL | 1626 | 16304 | 670 | 3284 |
| CI | mL/min/kg | 10.04 | - | 22.06 | - |
| Vdss | L/kg | 3.3 | - | 13.30 | 22.06 |
| Beta $t_{1/2}$ | hr | 4.76 | NC | 8.02 | NC |
| $C_{max}/C_0$ | ng/mL | 443.26 | 1026.00 | 96.2 | 184.3 |

I.V. PK: 1mg/kg
P.O. PK: 30mg/kg

**Fig. S2. (A)** Plasma concentrations over time of free AU-24118 in mice upon oral dosing (PO) of indicated concentrations of AU-24118. **(B)** Pharmacokinetics metrics for AU-24118 in mice and rats through intravenous injection and oral gavage. NC = not calculated, Cmax = Peak Concentration, C0 = Initial Concentration, $AUC_{0\text{-}last}$ = Area Under the Curve from Time 0 to Last Measurable Concentration, $AUC_{0\text{-}INF}$ = Area Under the Curve from Time 0 to Infinity, CI = Clearance, Vdss = Volume of Distribution at Steady State, Beta $t_{1/2}$ = Elimination Half-life, Tmax = Time to Peak Concentration.

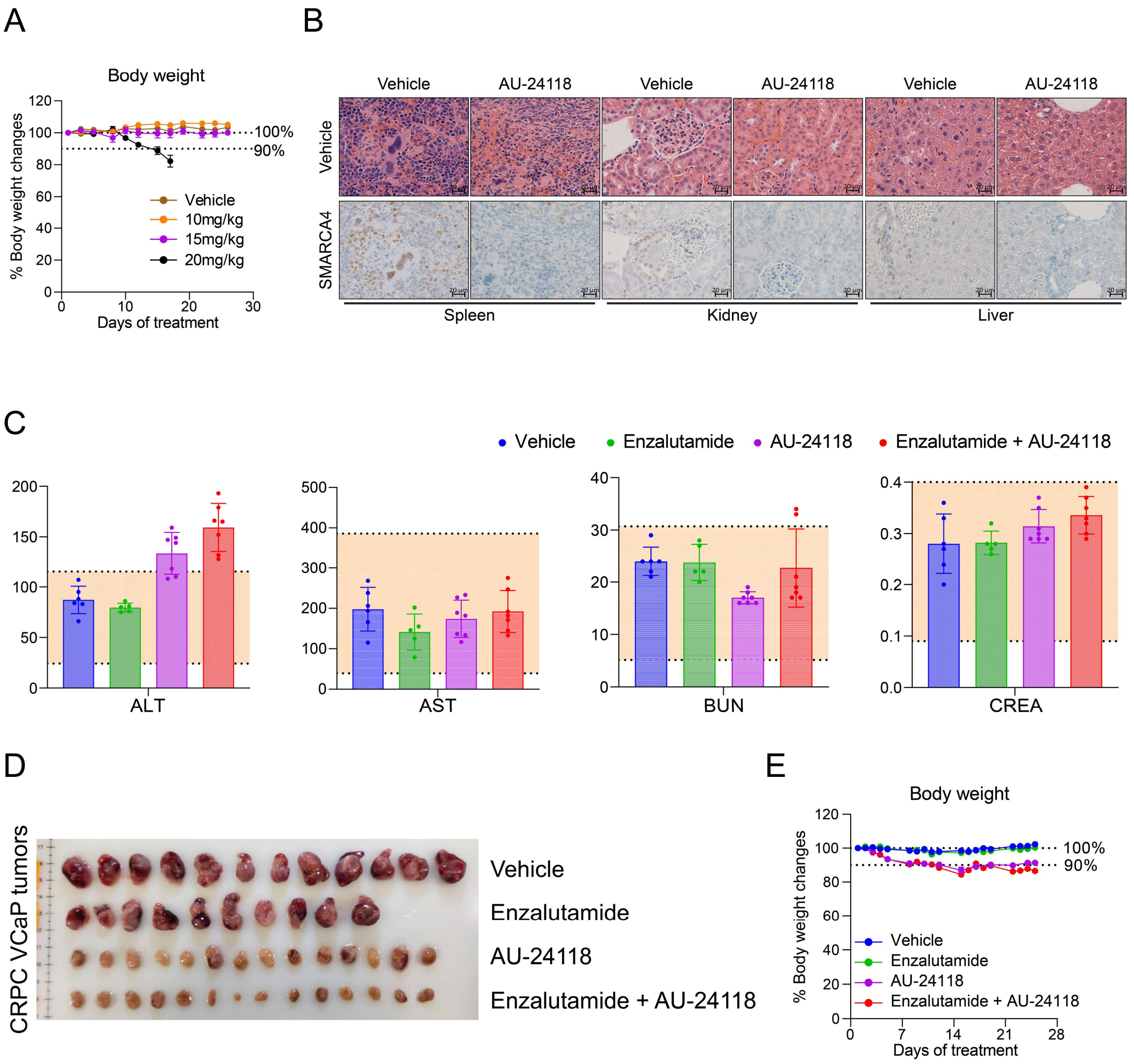


**Fig. S3. (A)** Percent body weight measurements of CB17 SCID mice throughout treatment course with indicated doses of vehicle or AU-24118. **(B)** Representative H&E staining images with corresponding IHC analyses for SMARCA4 in the indicated organs upon AU-24118 (15 mg/kg) treatment compared to vehicle controls (scale=20 μm). **(C)** Blood tests assessing liver and kidney function to evaluate toxicity of vehicle, enzalutamide (10 mg/kg), AU-24118 (15 mg/kg), and AU-24118 (15 mg/kg) + enzalutamide (10 mg/kg) after 25 days treatment in CB17 SCID mice. Normal range is indicated by orange shading. ALT = alanine aminotransferase, AST = aspartate aminotransferase, BUN = blood urea nitrogen, and CREA = creatinine. **(D)** Photographs of tumors from vehicle, enzalutamide, AU-24118, and AU-24118+enzalutamide groups from VCaP-CRPC study. (vehicle: n = 12, enzalutamide: n = 10, AU-24118: n = 14, AU-15330+enzalutamide: n = 14). **(E)** Relative (normalized to day 0) body weight measurements of CB17 SCID mice from the CRPC VCaP xenograft model throughout the treatment course with the indicated treatment.

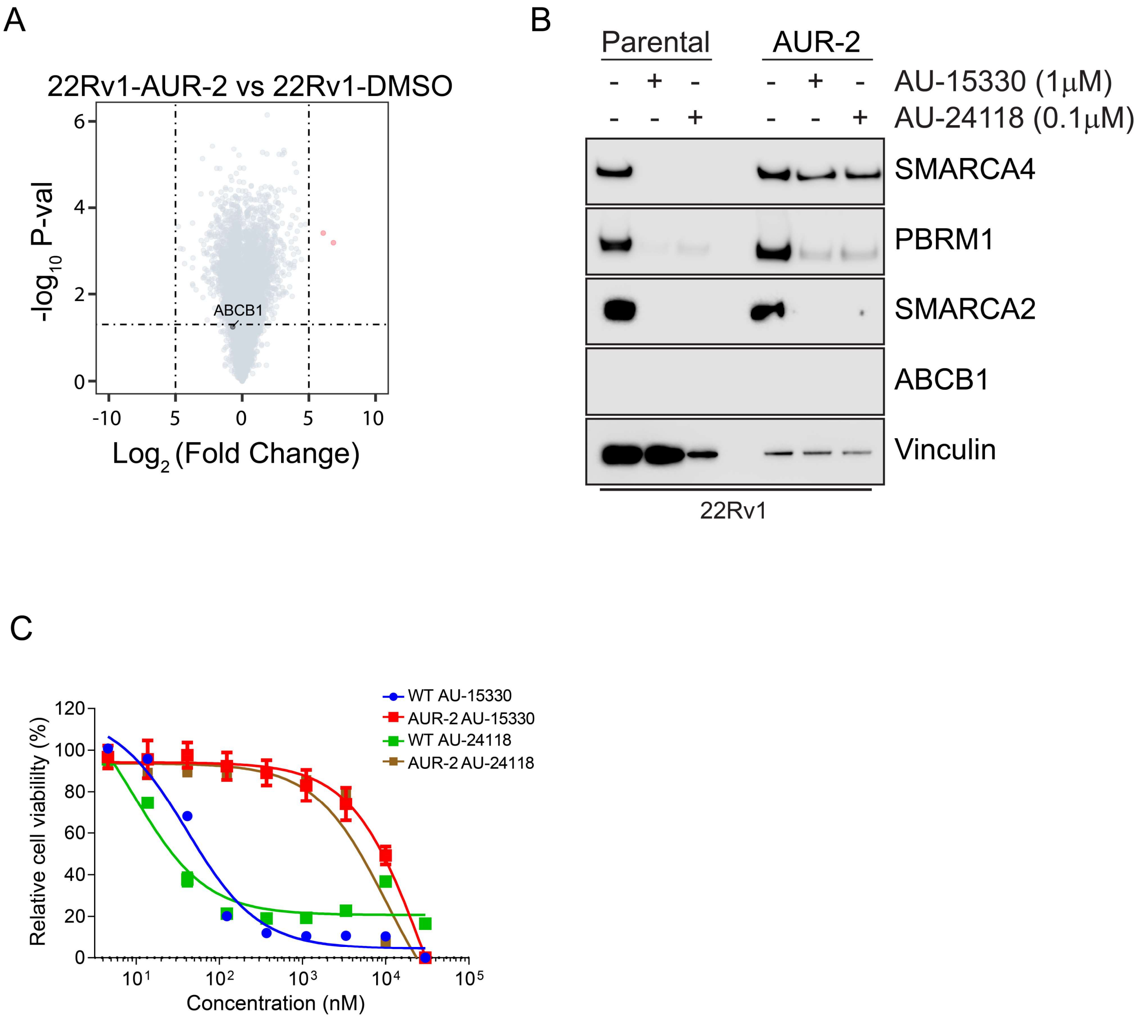


**Fig. S4. (A)** Volcano plot visualizing overall transcriptomic alterations as assessed by RNA-seq in 22Rv1-AUR-2 versus 22Rv1-WT cells. **(B)** Immunoblots of 22Rv1-wild type (WT) or 22Rv1 AUR-2 cells treated with 4h AU-15330 or AU-24118 with indicated concentration showing changes in the labeled targets. Vinculin was used as a loading control. This experiment was repeated independently twice. **(C)** Dose-response curves of 22Rv1-wild type (WT) or 22Rv1 AUR-2 cells treated with AU-15330 or AU-24118. Data are presented as mean +/− SD (n = 6) from one-of-three independent experiments.

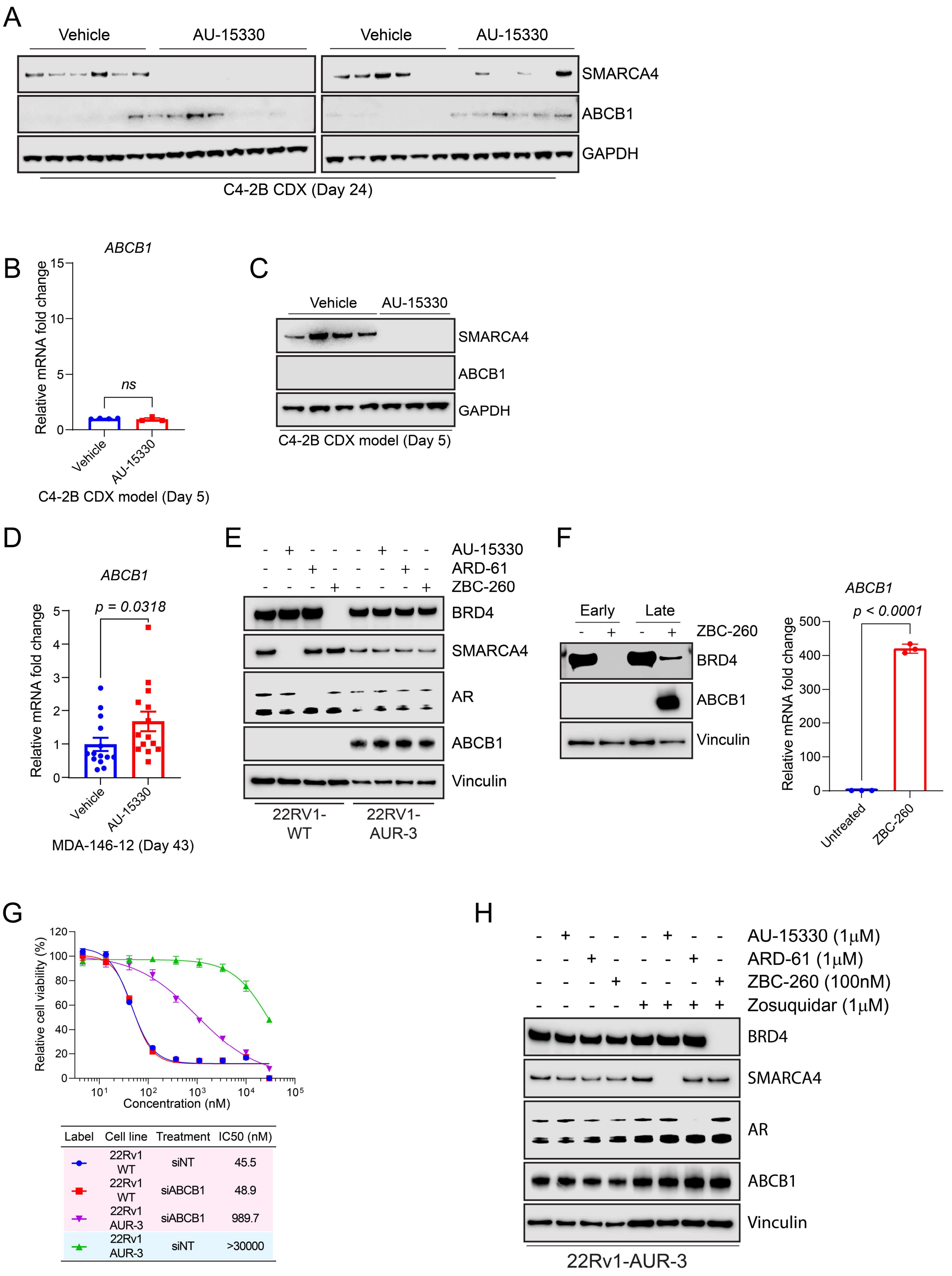


**Fig. S5. (A)** Immunoblot illustrating levels of ABCB1 proteins in C4-2B xenografts after long-term (24 days) AU-15330 treatment. Vinculin is utilized as the loading control across immunoblots. CDX, cell line-derived xenograft. **(B)** qPCR of C4-2B xenografts treated with AU-15330 or vehicle for 5 days showing changes in RNA levels of *ABCB1*. Data shown are technical triplicates. T-tests were performed as two-tailed t-tests assuming equal variances. **(C)** Immunoblot illustrating levels of ABCB1 in C4-2B xenografts after short-term (5 days) AU-15330 treatment. Vinculin is utilized as the loading control across immunoblots. **(D)** qPCR of MDA-146-12 xenografts after treatment with AU-15330 or vehicle for 43 days showing changes in RNA levels of *ABCB1*. Data shown are technical triplicates. T-tests were performed as two-tailed t-tests assuming equal variances. **(E)** Immunoblots of 22Rv1-wild type (WT) or 22Rv1 AUR-3 cells treated for 4 hours with 1 μM AU-15330, 1 μM ARD-61, or 100 nM ZBC-260 showing changes in the indicated protein levels. Vinculin was used as a loading control. This experiment was repeated independently twice. **(F)** (Left) Immunoblots of 22Rv1 cells treated with DMSO or ZBC-260 (0.1 μM) at an early (3 weeks) or late (4 weeks) time point showing changes in the labeled protein targets. Vinculin was utilized as the loading control. (Right) qPCR of 22Rv1 cells upon treatment with ZBC-260 (0.1 μM) for 4 weeks showing changes in *ABCB1* mRNA levels. T-tests were performed as two-tailed t-tests assuming equal variances. **(G)** (Top) Dose-response curves of 22Rv1-wild type (WT) or 22Rv1 AUR-3 cells treated with AU-15330 with siNT (Negative control) or siABCB1. Data are presented as mean +/− SD (n = 6) from one-of-three independent experiments. (Bottom) $IC_{50}$ values calculated from dose-response curve experiment. **(H)** Immunoblots of 22Rv1 AUR-3 cells treated with AU-15330 (1 μM), ARD-61 (1 μM), or ZBC-260 (100 nM) with or without zosuquidar (1 μM). Vinculin was used as a loading control. This experiment was repeated independently twice.

**Table S1.** Antibodies used in this study.

| Antigen | Vendor | Catalog Number | Application | Antibody Dilution |
|---|---|---|---|---|
| SMARCA2/BRM | Bethyl Laboratories | A301-016A | Western Blot | 1:1000 |
| SMARCA4/BRG1 | Cell Signaling Technology | 52251S | Western Blot | 1:1000 |
| PBRM1 | Bethyl Laboratories | A301-591A | Western Blot | 1:1000 |
| Vinculin | Cell Signaling Technology | 18799S | Western Blot | 1:1000 |
| c-Myc | Abcam | ab32072 | Western Blot | 1:1000 |
| Cleaved PARP (Asp214) | Cell Signaling Technology | 9541 | Western Blot | 1:1000 |
| AR | Millipore Sigma | 06-680 | Western Blot | 1:1000 |
| ERG | Abcam | ab92513 | Western Blot | 1:1000 |
| KLK3 | Cell Signaling Technology | 5365s | Western Blot | 1:1000 |
| FLAG | Millipore Sigma | F1804-200G | Western Blot | 1:1000 |
| ABCB1 | Cell Signaling Technology | 13342S | Western Blot | 1:1000 |
| Histone H3 | Cell Signaling Technology | 3638S | Western Blot | 1:1000 |
| BRD4 | Bethyl Laboratories | A700-004CF | Western Blot | 1:1000 |
| BRG1 | Abcam | ab108318 | IHC | Pre-dilute |
| AR | Abcam | ab133273 | IHC | 1:2000 |
| ERG | Ventana | 790-4576 | IHC | Pre-dilute |
| C-Myc | Ventana | 790-4628 | IHC | Pre-dilute |
| Ki-67 | Ventana | 790-4286 | IHC | Pre-dilute |

**Table S2.** Sequences of RT-qPCR primers used in this study.

| Primer | Sequence |
|---|---|
| ABCB1-forward | GCTGTCAAGGAAGCCAATGCCT |
| ABCB1-reverse | TGCAATGGCGATCCTCTGCTTC |
